# Click-ExM enables expansion microscopy for all biomolecules

**DOI:** 10.1101/2020.03.19.998039

**Authors:** De-en Sun, Xinqi Fan, Hao Zhang, Zhimin Huang, Qi Tang, Wei Li, Jinyi Bai, Xiaoguang Lei, Xing Chen

## Abstract

Expansion microscopy (ExM) allows super-resolution imaging on conventional fluorescence microscopes, but has been limited to proteins and nucleic acids. Here we develop click-ExM, which integrates click-labeling into ExM to enable a “one-stop-shop” method for nanoscale imaging of various types of biomolecules. Using 18 clickable labels for click-ExM imaging of DNA, RNA, proteins, lipids, glycans and small molecules, we demonstrate its universality, compatibility with signal-amplification techniques, and broad applications in cellular imaging.

---

By physically expanding proteins or RNA of fixed specimens embedded in a swellable polymer hydrogel, expansion microscopy (ExM) enables nanoscale imaging by using conventional diffraction-limited microscopes^1^. As a critical step of ExM, the biomolecules or the labeled fluorophores need to be covalently anchored into the polymer network and preserved during homogenization of the fixed cells (e.g., by strong protease digestion) to ensure isotropic expansion. This imposes a great challenge to develop tailored protocols for specific imaging targets including proteins and RNA; however, other types of biomolecules such as lipids, glycans, and small molecules are currently not amenable to ExM.

To expand the applicability of ExM, we developed click-ExM, a variant of ExM, into which a unified method to label, anchor, and preserve fluorescent signals for all kinds of biomolecules is integrated (**Fig. 1a**). To develop such a unified protocol, we exploited click-labeling, which has emerged as a versatile tool for fluorescence imaging of various biomolecules^2^. DNA, RNA, proteins, glycans and lipids can all be metabolically labeled with a biorthogonal or “clickable” functional group (e.g., the azide or alkyne), which is subsequently conjugated with fluorescent probes via biorthogonal chemistry such as Cu(I)-catalyzed azide-alkyne cycloaddition (CuAAC or click chemistry) and copper-free click chemistry (**Supplementary Fig. 1**). In click-ExM, the metabolically labeled biomolecules are reacted with azide-biotin or alkyne-biotin via click chemistry, followed by staining with fluorescently labeled streptavidin (**Fig. 1a** and **Supplementary Fig. 2**). The streptavidin-fluorophore conjugates serve as a tri-functional probe, which binds biotin, anchors into the gel, and presents fluorescence. Glutaraldehyde (GA) or the succinimidyl ester of 6- ((acryloyl)amino)hexanoic acid (AcX) could react with the lysine residues of streptavidin and provide the anchoring groups. Up to 14 acrylamide groups were conjugated onto streptavidin by using AcX as the anchoring agent, ensuring efficient incorporation during gelation (**Supplementary Fig. 3a,b**). More importantly, streptavidin was resistant to strong protease digestion (i.e., by proteinase K) (**Supplementary Fig. 3c**), consistent with the previously observed preservation of streptavidin in protein retention ExM^3^. These results indicate that streptavidin should be readily cross-linked into the gel and survive through homogenization and expansion.

**Fig. 1.**
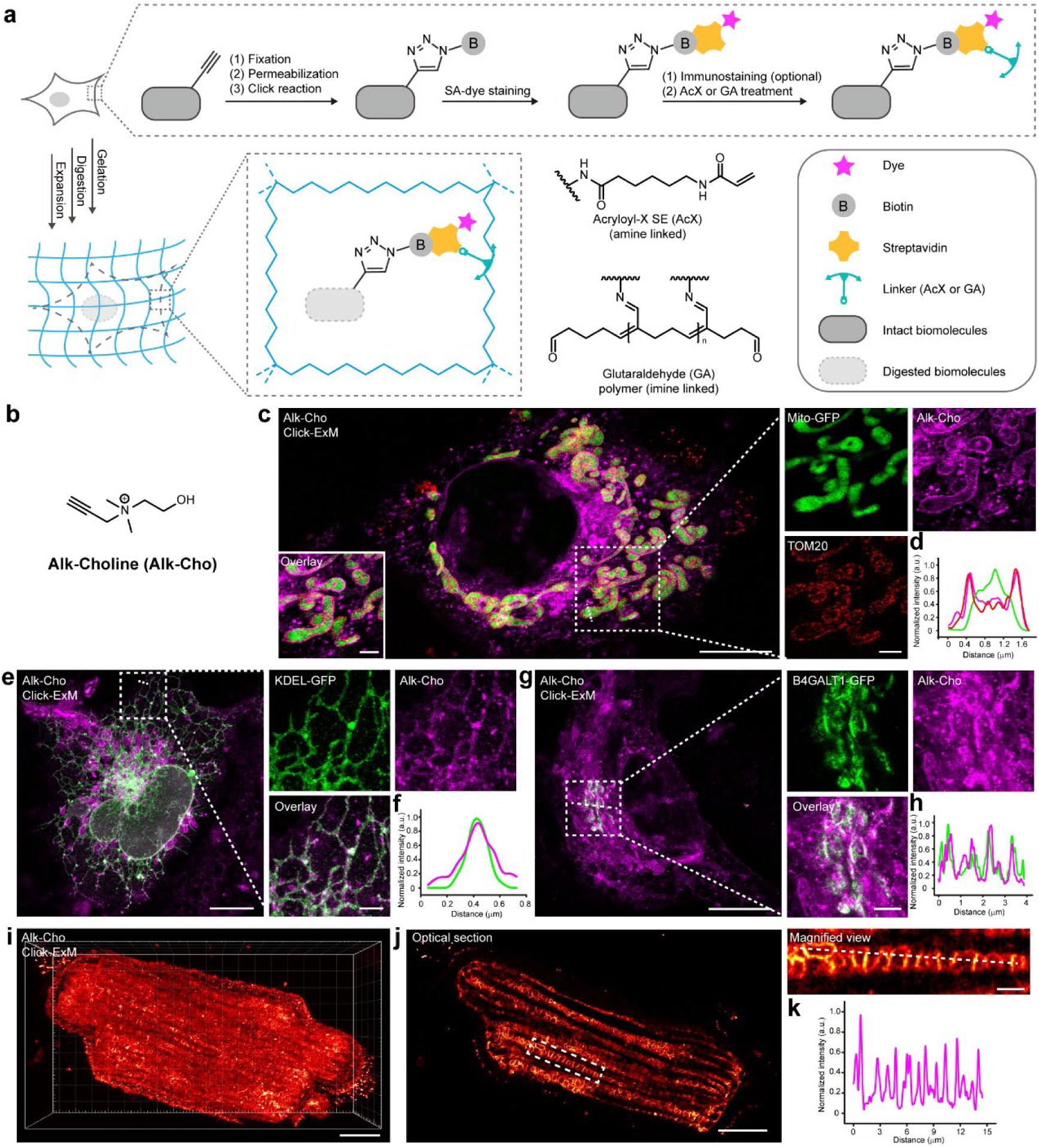
Click-ExM imaging of lipids. (**a**) Schematic of the click-ExM workflow. After metabolic labeling of biomolecules with alkynes, the cells are reacted with azide-biotin via click chemistry and stained with streptavidin-dye (SA-dye), followed with the standard ExM procedure. (**b**) Chemical structure of alk-Cho. (**c**) Click-ExM images of alk-Cho-labeled phospholipids (AF555, magenta) in COS-7 cells. Mito-GFP (green) was expressed in the mitochondrial matrix and TOM20 located in the mitochondrial outer membrane was immunostained with AF647 (red). (**d**) Fluorescence intensity profiles of mito-GFP (green), alk-Cho (magenta) and TOM20 (red) along the dotted line in the boxed region in **c**. a.u., arbitrary units. (**e**) Click-ExM images of alk-Cho-labeled phospholipids (AF555, magenta) and ER expressing KDEL-GFP (green) in COS-7 cells. (**f**) Fluorescence intensity profiles of KDEL-GFP (green) and alk-Cho (magenta) along the dotted line in **e**. (**g**) Click-ExM images of alk-Cho-labeled phospholipids (AF555, magenta) in COS-7 cells with the Golgi expressing B4GALT1-GFP (green). (**h**) Fluorescence intensity profiles of B4GALT1-GFP (green) and alk-Cho (magenta) along the dotted line in **g**. (**i**) 3D click-ExM image of alk-Cho-labeled phospholipids in rat cardiomyocyte (AF555, red hot). (**j**) Optical section and magnified image of **i**. (**k**) Fluorescence profile of alk-Cho (magenta) along the dotted line in **j**. Scale bars: 10 μm (**c, e, g, i, j**) and 2 μm (inset of **c, e, g, j**). All distances and scale bars are corresponding to pre-expansion dimension. GA was used for anchoring. All experiments were independently performed ≥3 times with a confocal microscope; representative data are shown.

We first demonstrated click-ExM for imaging lipids, a major biomacromolecule that has not been visualized by ExM. The protein-retention ExM protocol with an average expansion of 4.5-fold was adapted for this work^3,4^ (**Supplementary Fig. 4a,b**). COS-7 cells were treated with alkyne-choline (alk-Cho) to metabolically label Cho-containing phospholipids, the most abundant phospholipids in cellular membranes^5^ (**Fig. 1b**). After reacting the incorporated alk-Cho with azide-TAMRA, the membranes could be visualized by confocal microscopy before expansion, but the fluorescence was completely lost after the standard ExM procedure (**Supplementary Fig. 4c,d)**. Membrane staining confirmed that lipids could not be retained through the expansion process in ExM (**Supplementary Fig. 4e)**. We therefore applied click-ExM, in which the alk-Cho-treated cells were fixed and permealized with saponin, followed by click reaction with azide-biotin, staining with streptavidin-Alexa Fluor 555 (AF555), and AcX or GA treatment (**Fig. 1a**). After gelation, digestion, and expansion, the AF555 fluorescence was well preserved and the alk-Cho-incorporated membranes were visualized with super resolution (**Fig. 1b-k, Supplementary Fig. 4f** and **Supplementary Fig. 5a-f**). By co-staining of the mitochondrial markers and multi-color ExM, the distribution of Cho-containing phospholipids on the outer membrane of mitochondria was clearly observed (**Fig. 1c,d** and **Supplementary Fig. 5a**). Similarly, the structures of endoplasmic reticulum (ER) and Golgi apparatus could be resolved by click-ExM (**Fig. 1e-h** and **Supplementary Fig. 5b,c**). We then applied click-ExM to rat cardiomyocytes treated with alk-Cho, in which the network of interconnected and highly dense mitochondria was visualized (**Fig. 1i-k** and **Supplementary Fig. 5d-f**). Various lipid reporters including alkyne-palmitic acid, alkyne-farnesol, alkyne-myristic acid, alkyne-stearatic acid, and azide-cholesterol were compatible with click-ExM, which enabled super-resolution imaging of the distribution of fatty acids, prenol lipids and sterol lipids in COS-7 cells (**Supplementary Fig. 5g**).

In addition, protein carbonylation by 4-hydroxy-2-nonenal (HNE), a lipid-derived electrophile that reacts with cysteine residues, was visualized by click-ExM by conjugating the resulting aldehyde group on proteins with aminooxy-alkyne^6^ (**Supplementary Fig. 5h**). These results demonstrate the broad utility of click-ExM in lipid imaging.

The volumetric dilution of fluorophores in ExM reduces the signal intensity, a disadvantage which can be overcome by signal-amplification techniques. Because amplification by the immunosignal hybridization chain reaction (isHCR) has been demonstrated on streptavidin with the biotin-DNA HCR initiator conjugate^7^, click-ExM should be compatible with isHCR (**Supplementary Fig. 6a**). Furthermore, we developed a controllable signal amplification protocol by exploiting the multivalence of streptavidin (**Supplementary Fig. 6b**). By synthesizing a biotin trimer (**Supplementary Note 1**), streptavidin staining could be iteratively performed with the signal gradually increased in each round of amplification (**Supplementary Fig. 6c**).

Next, we applied click-ExM for imaging glycans, another major biomacromolecule not yet amenable to ExM. The sialoglycans on the membrane of HeLa cells were metabolically labeled with azido sialic acid (SiaNAz; **Fig. 2a**). Click-ExM revealed that the SiaNAz-labeled glycans were widely distributed on the basal plasma membrane (**Fig. 2b**, left). At super resolution, the sialoglycans on the hollow membrane protrusions such as microvilli were clearly resolved (**Fig. 2b**, right). These observations were consistent with earlier studies on sialoglycans using localization-based super-resolution microscopy^8-10^. Click-ExM imaging of sialoglycans was then performed on rat hippocampal neurons treated with *N*-azidoacetylmannosamine (ManNAz; **Fig. 2a**), the metabolic precursor of SiaNAz. With click-ExM, the tube-like structure of neurites with an inner-diameter of ∼500 nm could be clearly delineated by sialoglycans on the membrane (**Fig. 2c**, top). By contrast, the tubular structure was unresolvable before expansion (**Fig. 2c**, bottom). In rat cardiomyocytes which had a relatively low labeling with ManNAz, click-ExM with isHCR amplification revealed sialoglycans on the orderly spaced transverse tubule (T-tubule) network (**Fig. 2d**, top), with a much better resolution than conventional confocal microscopy^11^ (**Fig. 2d**, bottom and **Supplementary Fig. 7a**).

**Fig. 2.**
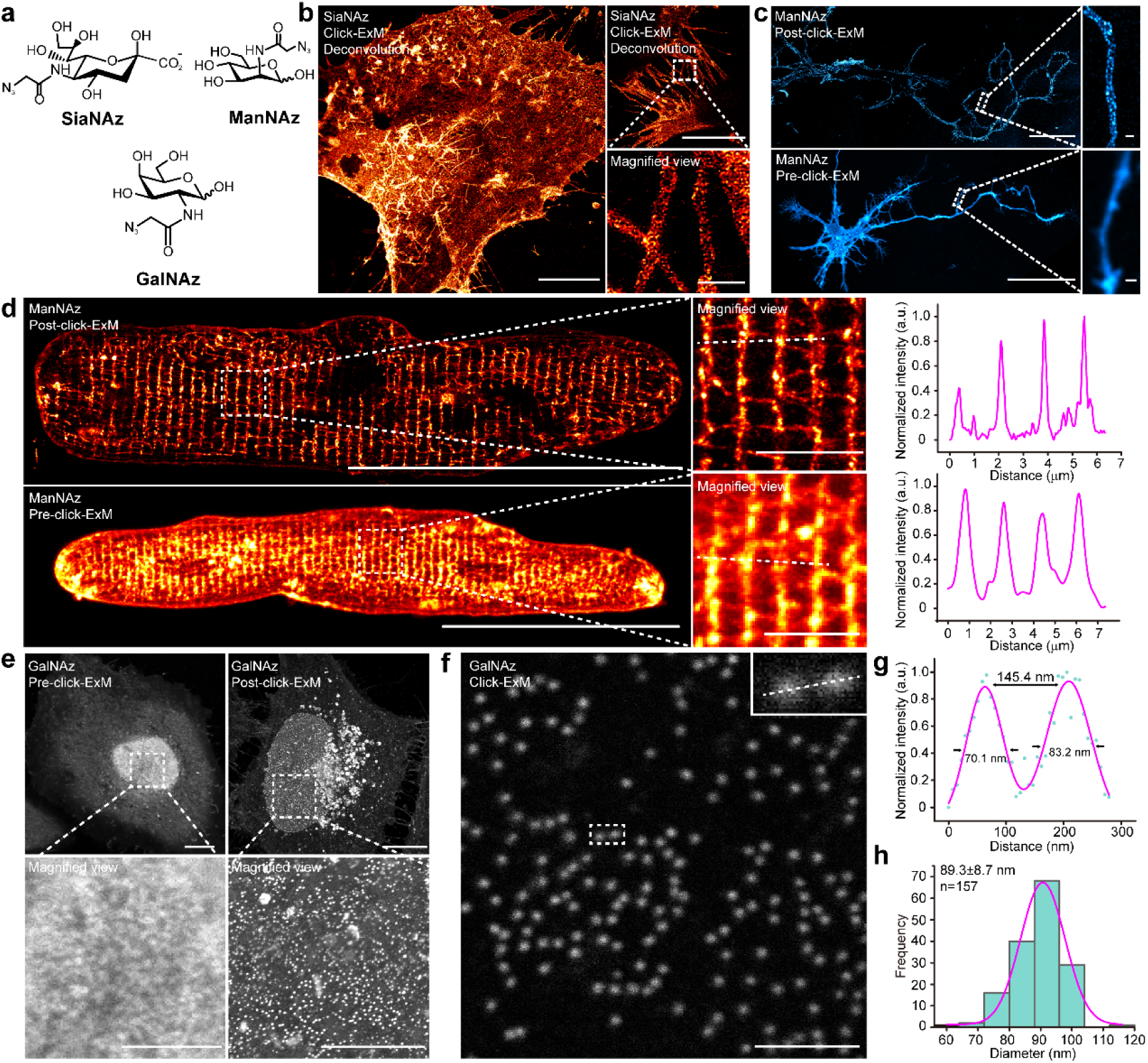
Click-ExM imaging of glycans. (**a**) Chemical structures of SiaNAz, ManNAz and GalNAz. (**b**) Click-ExM images of sialoglycans (AF555, red hot) in SiaNAz-treated HeLa cells after deconvolution. (**c**) Pre- and post-click-ExM images of sialoglycans (AF488, cyan hot) in ManNAz-treated rat hippocampus neurons. (**d**) Pre- and post-click-ExM images of sialoglycans (AF546, red hot) in ManNAz-treated rat cardiomyocytes after isHCR amplification. Fluorescence profiles along the dotted lines in the boxed regions from pre- and post-click-ExM images were shown. (**e**) Pre- and post-click-ExM images of O-GlcNAc (AF555, gray) in GalNAz-treated HeLa cells. (**f**) Representative click-ExM image showing morphological details of GalNAz-labeled nucleoporins (AF488, gray) in HeLa cells. (**g**) Fluorescence profile of the boxed region in **f**. (**h**) Diameter distribution of GalNAz-labeled nucleoporins in **f**. Scale bars: 50 μm (**c, d**), 10 μm (**b, e**), 5 μm (inset of **d, e**) and 1 μm (inset of **b, c** and **f**). All distances and scale bars are corresponding to the pre-expansion dimension. AcX (**b, e, f**) or GA (**c, d**) was used for anchoring. All experiments were independently performed ≥3 times with a confocal microscope; representative data are shown.

We then applied click-ExM for imaging O-GlcNAcylation, modification of serine and threonine residues of various intracellular proteins with an *N*-acetylglucosamine (GlcNAc) monosaccharide. HeLa cells were treated with *N*-azidoacetylgalactosamine (GalNAz; **Fig. 2a**), which metabolically labeled both cell-surface glycans and O-GlcNAc^12^. After click reaction with alkyne-biotin, staining with streptavidin-AF555, fluorescence distributed in the cytoplasm and nucleus before and after expansion, presumably from the labeled O-GlcNAc (**Fig. 2e** and **Supplementary Fig. 7b**). Nucleoporin, the constituent building blocks of the nuclear pore complex (NPC), are heavily O-GlcNAcylated and can be stained by the O-GlcNAc-recognizing lectin wheat germ agglutinin (WGA), which have been utilized for super-resolution imaging of the central channel of NPC^13^. Heavy GalNAz labeling was co-localized with NPC staining on the nuclear envelop (**Supplementary Fig. 7c**). Using click-ExM, individual NPC was resolved with a diameter of ∼89 nm by metabolic O-GlcNAc labeling (**Fig. 2f-h** and **Supplementary Fig. 7d**). Click-ExM imaging of O-GlcNAc could also be performed by using the chemoenzymatic method based on a mutant galactosyltransferase (Y289L GalT) to label O-GlcNAc with GalNAz^14^ (**Supplementary Fig. 7e**). In addition, click-ExM in combination with chemoenzymatic labeling was generally applicable for nanoscale imaging of other glycans, such as the cancer-associated *N*-acetyllactosamine (LacNAc) on the surface of HeLa cells^15^ (**Supplementary Fig. 7f**). Of note, the LacNAc level was relatively low and the signal was amplified by the iterative biotin-streptavidin staining method.

To demonstrate the applicability of click-ExM for protein imaging, rat hippocampal neurons were treated with azidohomoalanine (AHA; **Fig. 3a**), a noncanonical amino acid serving as a surrogate for methionine, to metabolically label nascent proteins^16^. Click-ExM imaging revealed the distribution of newly synthesized proteins in somata and dendrites with super resolution (**Fig. 3b**).

**Fig. 3.**
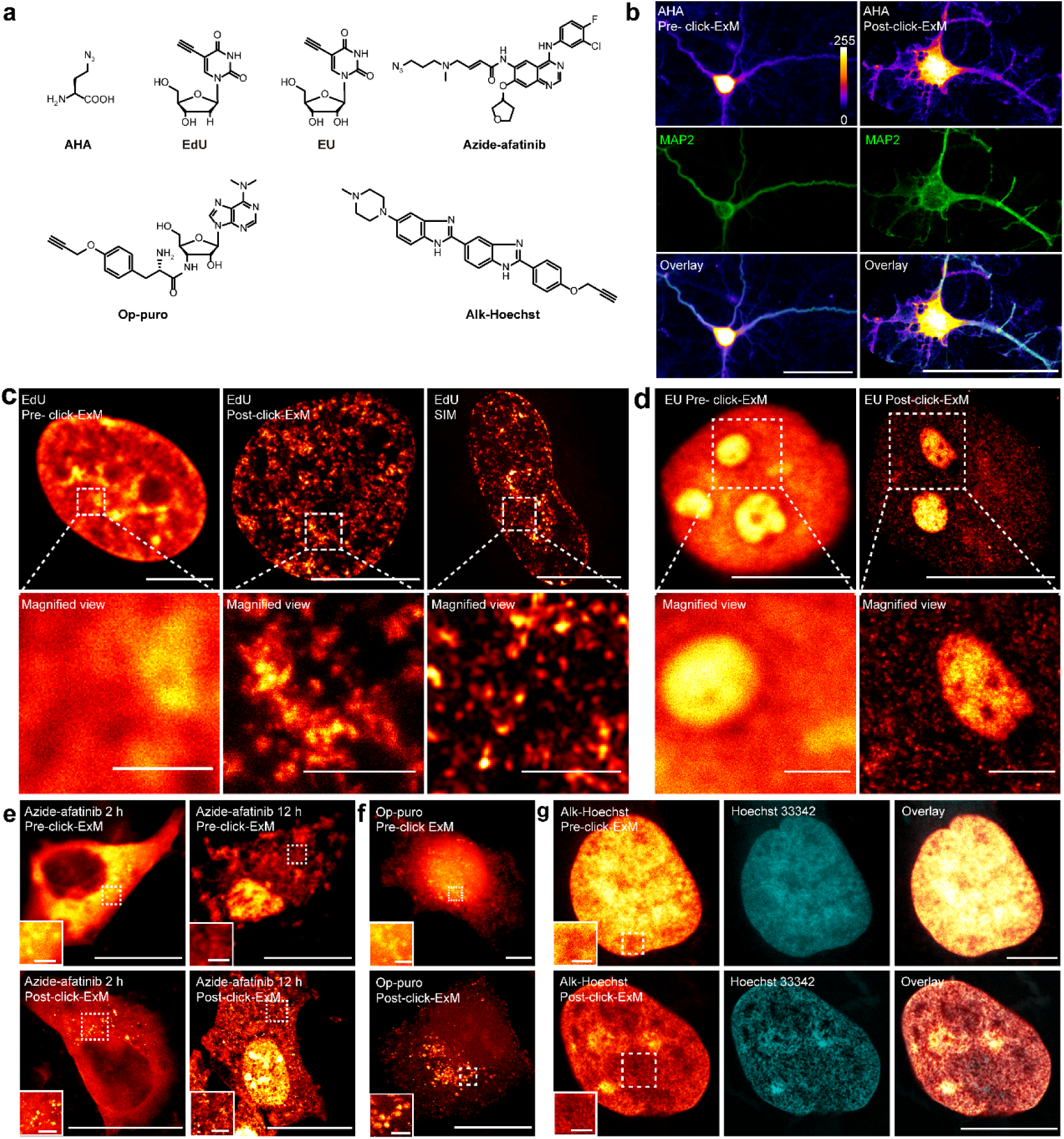
Click-ExM imaging of proteins, nucleic acids and small molecules. (**a**) Chemical structures of AHA, EdU, EU, azide-afatinib, O-propargyl-puromycin (Op-puro) and alk-Hoechst. (**b**) Pre- and post-click-ExM images of nascent proteins in AHA-treated rat hippocampus neurons. Color lookup table indicates fluorescence intensity (AF555, pixel intensities 0–255). MAP2 (AF488, green) was immunostained. (**c**) Pre- and post-click-ExM and SIM images of nascent DNA (AF555, red hot) in EdU- treated U2OS cells. (**d**) Pre- and post-click-ExM images of nascent RNA (AF555, red hot) in EU-treated HeLa cells. (**e**) Pre- and post-click-ExM images of HeLa cells treated with azide-afatinib (AF555, red hot) for 2 h and 12 h. (**f**) Pre- and post-click-ExM images of nascent peptides (AF555, red hot) in Op-puro-treated HeLa cells. (**g**) Pre- and post-click-ExM images of DNA (AF555, red hot) in alk-Hoechst-labeled U2OS cells. Nucleus were stained with Hoechst 33342 (cyan). Scale bars: 50 μm (**b**), 20 μm (**e**-**g**), 10 μm (**c, d**) and 2 μm (inset of **c**-**g**). Scale bars are corresponding to the pre-expansion dimension. AcX was used for anchoring. All experiments were independently performed ≥3 times with a confocal microscope except the SIM imaging (two independent experiments); representative data are shown.

For nucleic acid imaging, we used 5-ethynyl-2’-deoxyuridine (EdU; **Fig. 3a**) to metabolically label nascent DNA^17^. Under click-ExM, the fine structure of chromatin in cell nuclei was successfully resolved at a resolution comparable to structured illumination microscopy (**Fig. 3c**). With super resolution, the distribution of heterochromatin and euchromatin was differentiated. During S phase, EdU labeling and click-ExM clearly revealed the boundaries of replication domains, which are co-replicating Mb-sized segments of chromosomal DNA (**Supplementary Fig. 8a**). By using multi-color click-ExM, we showed that the newly synthesized DNA was largely co-localized with histone H3 (**Supplementary Fig. 8b**). In addition, nascent RNA in HeLa cells was metabolically incorporated with 5-ethylnyluridine (EU) and visualized by click-ExM^18^ (**Fig. 3d**). In the nucleus, intense labeling in nucleoli, presumably from pre-rRNA, and diffused labeling in the nucleoplasm, presumably from mRNA, were observed at nanoscale resolution.

In addition to biomacromolecules, we envisioned that click-ExM would enable super-resolution imaging of small molecules, including drugs, metabolites, and chemical probes, in the cells. The small molecules should tolerate modification with an azide or alkyne without interfering with their function and cellular localization. Afatinib is an irreversible inhibitor for ErbB family and has been approved by FDA to treat metastatic non-small cell lung cancer with non-resistant epidermal growth factor receptor (EGFR) mutations^19^. We installed an azide at the position so that the activity of afatinib was not affected (**Fig. 3a**). Click-ExM revealed the distribution of azide-afatinib in HeLa cells, and plasma membrane bledding during afatinib-induced cell apoptosis was observed (**Fig. 3b**). Puromycin is an aminonucleoside antibiotic that terminates protein synthesis by covalent conjugation with nascent polypeptide chains. Here, an alkyne-containing puromycin analog, O-propargyl-puromycin^20^ (OP-puro; **Fig. 3a**), was used for click-ExM. Intense labeling was observed in various cytoplasmic foci, which probably resulted from rapid degradation of misfolded peptides by proteasome (**Fig. 3f**). The widely used DNA stain Hoechst 33258 was derivatized with an alkyne (**Fig. 3a**), with which the nuclei were visualized with AF555 by click-ExM (**Fig. 3g**).

ExM has emerged as an attractive approach to perform super-resolution imaging by using conventional microscopes. Click-ExM developed in this work provides a unified and convenient protocol for nanoscale imaging of essentially all biomolecules, thus expanding the applicability of ExM to molecules not being imaged before (e.g., lipids). Click-ExM exploits the click-labeling method, which is applicable for various biomolecules and is advancing rapidly. By converting the click labels to streptavidin-dye, click-ExM adapts the well-established ExM workflow. Furthermore, click-ExM is compatible with various signal amplification techniques. A promising future direction is to integrate different click reactions to enable visualization of multiple kinds of biomolecules in the same sample (**Supplementary Fig. 9**).

## Acknowledgements

We thank Prof. Shiqiang Wang and Xiaoting Wang for providing rat cardiomyocytes, Prof. Yujie Sun for providing COS-7 cells, Prof. Liangyi Chen for providing the plasmids of KDEL-GFP, and B4GALT1-GFP, Prof. Peng Zou for providing the mito-GFP plasmid, Prof. Chu Wang for providing AOyne, Prof. Baoliang Song for providing azide-cholesterol, Dr. Wen Zhou (Mass Spectrometry Facilities of the National Center for Protein Sciences, Peking University) for the help on MALDI mass experiment, and Dr. Chunyan Shan (Core Facilities at School of Life Sciences, Peking University) for the help on SIM imaging. This work is supported by the National Key Research and Development Projects (No. 2018YFA0507600) and the National Natural Science Foundation of China (No. 91753206 and No. 21521003).

## Author contributions

D.S. and X.C. conceived the project; D.S. performed experiments with the help of X.F., H.Z., Z.H., Q.T., W.L., J.B., and X.L.; D.S. and X.C. analyzed the data and wrote the manuscript.

## ONLINE METHODS

### Regents and antibodies

All chemical reagents were obtained from commercial suppliers, and used without further purification unless otherwise noted. Copper (II) sulfate pentahydrate (C8027), (+)-sodium L-ascorbate (A4034), glutaraldehyde (G7651), acrylamide (A9099), N,N′-methylenebisacrylamide (M7279), sodium acrylate (408220), N,N,N,N - tetramethylethylenediamine (TEMED) (T7024), ammonium persulfate (A3678), dextran sulfate (D8906), biotin (14400), tris[(1-hydroxypropyl-1H-1,2,3-triazol-4-yl)methyl]amine (THPTA) (762342) and Hoechest 33342 (B2261) were purchased from Sigma-Aldrich. 5-ethynyl-2 -deoxyuridine (EdU) (HR-010908) and 5-ethynyluridine (EU) (HR-010907) were purchased from Wuhu Huaren Science and Technology. GalNAz (SAM703) was purchased from Jinan Samuel Pharmaceutical Co., Ltd. Azide-TAMRA (AZ109), biotin-PEG_3_-amine (1187), alkyne-myristic acid (1164), alkyne-palmitic acid (1165), alkyne-stearic acid (1166), DBCO-biotin (A116), 2-[4-{(bis[(1-tert-butyl-1H-1,2,3-triazol-4-yl)methyl]amino)methyl}-1H-1,2,3-triazol-1-yl]acetic acid (BTTAA) (1236), azide-PEG_3_-biotin (AZ104), and alkyne-PEG_4_-biotin (TA105) were purchased from Click Chemistry Tools. Azide-PEG_3_-FLAG (CLK-032) was purchased from Jena Bioscience. Acryloyl-X (AcX) (A20770), streptavidin (S888), streptavidin Alexa Fluor 488 (S32354) and streptavidin Alexa Fluor 555 (S32355) were purchased from Thermo Scientific. Azidohomoalanine (AHA) (A584003), saponin (482689) and Hoechst 33258 (398607) were purchased from J&K Scientific. Proteinase K (P8107S) and DNase I (M0303S) were purchased from NEB.

For immunostaining, the following primary and secondary antibodies were used in this study: anti-α-tublin (Abcam, ab52866, 1:250), anti-TOM20 (Abcam, ab186734, 1:250), Anti-O-Linked *N*-Acetylglucosamine (RL2) (Abcam, ab2739, 1:100), anti-NUP133 (Abcam, ab155990, 1:100), anti-caveolin 3 (Santa Cruz, sc-5310, 1:50), anti-MAP2 (Abcam, ab32454, 1:200), anti-Histone H3 (Abcam, ab176842, 1:2,000), Anti-DDDDK tag (Abcam, ab205606, 1:100), goat anti-mouse secondary antibody, Alexa Fluor 488 (Thermo, A-11001, 1:500), goat anti-mouse secondary antibody, Alexa Fluor 546 (Thermo, A-11003, 1:500), goat anti-rabbit secondary antibody, Alexa Fluor 488 (Thermo, A-11034, 1:200), goat anti-rabbit secondary antibody, Alexa Fluor 555 (Thermo, A-21429, 1:500), goat anti-rabbit secondary antibody, Alexa Fluor 647 (Thermo, A-21245, 1:1,000) and biotinylated goat anti-rabbit secondary antibody (Proteintech, SA00004-2, 1:200).

### Compound synthesis

Alkyne-choline^5^, O-propargyl-puromycin^20^, alkyne-farnesol^21^, SiaNAz^22^ and ManNAz^23^ were synthesized as previously described. The synthesis of biotin trimer, alkyne-Hoechst and azide-afatinib is described in **Supplementary Note 1** in the Supplementary Information.

### Cell culture

HeLa (ATCC, CCL-2), CHO (ATCC, CCL-61), U2OS (3111C0001CCC000028, National Infrastructure of Cell Line Resource, Beijing, China) and COS-7 (Cell Bank, Chinese Academy of Science, Shanghai, China) cells were cultured in Dulbecco’s modified Eagle’s medium (DMEM) supplemented with 10% (v/v) fetal bovine serum, 100 unit mL^-1^ penicillin, and 100 μg mL^-1^ streptomycin at 37°C in 5% CO_2_ atmosphere. Cells were used with passage number below 20 and free of mycoplasma contamination.

### Animals

Wild-type Sprague-Dawley rats were purchased from Vital River Laboratory Animal Center (Beijing, China) and kept under specific-pathogen-free (SPF) condition. All animal experiments were performed in accordance with guidelines approved by the Institutional Animal Care and Use Committee of Peking University accredited by AAALAC International. Single ventricular myocytes from adult Sprague-Dawley rats were enzymatically isolated by using Langendorff perfusion apparatus. Hippocampal neurons were dissociated from postnatal day 0 Sprague-Dawley rat pups. Briefly, hippocampi were dissected and triturated after incubating with 0.25% (w/v) trypsin-EDTA at 37°C for 14 min. Dissociated neurons were plated onto poly-D-lysine coated petri dishes and cultured in Neurobasal A medium supplemented with B-27 and Glutamax at 37°C for 7 d.

### Stoichiometric quantification of streptavidin-AcX reaction

10 μL of 1 mg mL^-1^ streptavidin was incubated with 2 μL of 10 mg mL^-1^ AcX in PBS overnight at room temperature (r.t.), and the molecular weight change of resulting reaction mixture was determined by MALDI-TOF/TOF Mass Spectrometer (AB Sciex 5800).

### Proteinase K sensitivity of streptavidin

Streptavidin (100 μg) was incubated with proteinase K (8 unit mL^-1^) in the presence or absence of biotin (600 μM) in digestion buffer (50 mM Tris pH 8.0, 1 mM EDTA, 0.1% (v/v) Triton X-100, 0.8 M guanidine HCl) at 37°C for 0 h, 2 h, 4 h, or overnight at r.t., followed by precipitation with 150 μL methanol, 37.5 μL chloroform and 100 μL water. The mixture was centrifuged at 18,000 g for 5 min for removal of the aqueous phase, and 100 μL methanol was added. After centrifugation and removal of methanol, the precipitate was re-suspended in 50 μL 1% (w/v) SDS, resolved on 12% SDS/PAGE and stained by Commassie Brilliant Blue (CBB).

### Immunostaining

Cells were fixed with 3% (w/v) formaldehyde and 0.1% (v/v) glutaraldehyde in PBS for 15 min and reduced with 0.1% (w/v) sodium borohydride for 7 min. After washing three times with 100 mM glycine in PBS, cells were permeabilized with 0.1% (v/v) Triton X-100 in PBS for 15 min. Cells were blocked in blocking buffer (1× PBS/5% (w/v) BSA/0.1% (v/v) Tween-20) for 30 min, and incubated with primary antibody in antibody dilution buffer (1× PBS/1% (w/v) BSA/0.1% (v/v) Tween-20) overnight at 4°C, then cells were washed three times with PBS and incubated with fluorophore-conjugated secondary antibody in antibody dilution buffer for 1 h and washed three times with PBS. Specifically, for α-tublin immunostaining, a cytoskeleton extraction step was performed before fixing the cells. Briefly the cells were extracted in cytoskeleton extraction buffer (0.2% (v/v) Triton X-100, 0.1 M PIPES, 1 mM EGTA, 1 mM MgCl_2_, pH 7.0) for 1 min at r.t. For TOM20 immunostaining in **Fig.1c** and **Supplementary Fig. 5a**, COS-7 cells were incubated with primary antibody in antibody dilution buffer (1× PBS/1% (w/v) BSA/0.1% (w/v) saponin) before ExM procedure, and AF647-conjugated secondary antibody was then incubated with gel overnight in antibody dilution buffer at 4°C after gelation and digestion step.

### Metabolic labeling of lipids

COS-7 cells were labeled with the following conditions: 100 μM alkyne-choline for 12 h, 200 μM alkyne-palmitic acid for 24 h, 200 μM alkyne-myristic acid for 24 h, 200 μM alkyne-stearic acid for 12 h, 100 μM alkyne-farnesol for 24 h, 20 μg mL^-1^ azide-cholesterol for 24 h (with 1 μM lovastatin). For HNE labeling, cells were incubated with 50 μM HNE in DMEM for 1 h, and after two washes with warm DMEM for 30 min, cells were incubated with 1 mM AOyne for 30 min. All the cells were fixed with 3% (w/v) formaldehyde and 0.1% (v/v) glutaraldehyde in PBS for 15 min and then reduced with 0.1% (w/v) sodium borohydride for 7 min. After washing three times with 100 mM glycine in PBS, cells were permeabilized with 0.1% (w/v) saponin in PBS for 5 min and washed three times with PBS. Then cells were incubated with azide-PEG_3_-biotin (50 μM), BTTAA-CuSO_4_ complex (50 μM CuSO_4_, BTTAA/CuSO_4_ 6:1, m/m) and sodium ascorbate (2.5 mM) in PBS at r.t. for 1 h, followed by five washes with PBS. For azide-cholesterol-labeled cells, permeabilization step with saponin is omitted and alkyne-PEG_4_-biotin was used for click-labeling. Cells were then incubated with 5 μg mL^-1^ streptavidin-dye in PBS containing 1% (w/v) BSA for 1 h, and washed three times with PBS. Lipid-labeled samples should be immediately executed into next steps to avoid any potential signal loss or distortion.

### Metabolic labeling of glycans

For O-GlcNAc labeling, HeLa and U2OS cells were incubated with 1 mM GalNAz for 48 h. Cells were fixed with 4% (w/v) formaldehyde in PBS for 15 min, washed three times with PBS, and permeabilized with 0.1% (v/v) Triton X-100 in PBS for 15 min. After three washes with PBS, cells were incubated with alkyne-PEG_4_-biotin (50 μM), BTTAA-CuSO_4_ complex (50 μM CuSO_4_, BTTAA/CuSO_4_ 6:1, m/m) and sodium ascorbate (2.5 mM) in PBS at r.t. for 1 h, followed by five washes with 0.1% (v/v) Tween-20 in PBS. For sialic acid labeling, 2 mM SiaNAz or ManNAz were used to treat HeLa cells, rat hippocampus neurons and rat cardiomyocytes for 48 h. After three washes with warm medium, cells were incubated with 100 μM DBCO-biotin in medium at 37 °C for 30 min and then washed with warm medium for 15 min. Cells were fixed with 3% (w/v) formaldehyde and 0.1% (v/v) glutaraldehyde in PBS for 15 min and reduced with 0.1% (w/v) sodium borohydride for 7 min. After washing three times with 100 mM glycine in PBS, Cells were permeabilized with 0.1% (w/v) saponin in PBS for 5 min and wash three times for subsequent immunostaining. Otherwise, permeabilization step is not necessary. All the samples were incubated with 5 μg mL^-1^ streptavidin-dye in PBS containing 1% (w/v) BSA for 1 h and washed three times with PBS. isHCR was used for signal amplification in ManNAz-labeled rat cardiomyocytes.

### Metabolic labeling of nucleic acids

For DNA labeling, U2OS cells were labeled with 10 μM EdU for 24 h. For RNA labeling, HeLa cells were labeled with 1 mM EU for 4 h. All the cells were fixed with 4% formaldehyde in PBS for 15 min. After three washes with PBS, cells were permeabilized with 0.1% (v/v) Triton X-100 in PBS for 15 min. Cells were incubated with azide-PEG_3_-biotin (50 μM), CuSO_4_ (50 μM), THPTA (200 μM) and sodium ascorbate (2.5 mM) in PBS, at r.t. for 1 h, followed by five washes with PBS. Cells were incubated with 5 μg mL^-1^ streptavidin-dye in PBS containing 1% (w/v) BSA for 1 h, and washed three times with PBS. Before gelation, EdU-labeled cells were treated with 30 unit mL^-1^ DNase I for 30 min at 37°C to fragment the genomic DNA.

### Chemoenzymatic labeling of glycans

For O-GlcNAc labeling, CHO cells were fixed, permeabilized with the same procedures of GalNAz labeling experiments, and then incubated in labeling solution containing 25 μg mL^-1^ Y289L GalT1, 500 μM UDP-GalNAz, 1 mM MnCl_2_, 50 mM NaCl, 20 mM HEPES (pH 7.9) and 2% (v/v) Nonidet P40 at 4 °C for 20 h. After five washes with PBS, cells were labeled with the same procedures of GalNAz labeling experiments. For LacNAc labeling, HeLa cells were washed three times with PBS, and incubated in labeling solution containing 20 mM MgCl_2_, 3 mM HEPES, 1% (v/v) FBS, 500 μM GDP-FucNAz and 50 μg mL^-1^ α 1,3-FucT at 37 °C for 15 min. After three washes with PBS, cells were labeled and fixed with the same procedures of SiaNAz or ManNAz labeling experiments. Iterative biotin trimer-streptavidin staining was used for signal amplification.

### Metabolic labeling of proteins

Rat hippocampus neurons were incubated with 1 mM AHA for 2 h. After three washes with PBS, neurons were fixed, permeabilized and labeled with the same procedures of GalNAz labeling experiments.

### Small molecules labeling

HeLa cells were treated with 100 μM azide-afatinib in DMEM for either 2 h or 12 h. After three washes with PBS, cells were fixed and permeabilized with the same procedures of lipids labeling experiments, then cells were incubated with alkyne-PEG_4_-biotin (50 μM), BTTAA-CuSO_4_ complex (50 μM CuSO_4_, BTTAA/CuSO_4_ 6:1, m/m) and sodium ascorbate (2.5 mM) in PBS at r.t. for 1 h, followed by five washes with PBS. The cells were incubated with 5 μg mL^-1^ streptavidin-dye in PBS containing 1% (w/v) BSA for 1 h and washed three times with PBS.

For Op-puro, HeLa cells were treated with 50 μM Op-puro in DMEM for 1 h. After washing with warm DMEM for 15 min, cells were fixed, permeabilized with the same procedures of GalNAz labeling experiments. Then cells were incubated with azide-PEG_3_-biotin (50 μM), BTTAA-CuSO_4_ complex (50 μM CuSO_4_, BTTAA/CuSO_4_ 6:1, m/m) and sodium ascorbate (2.5 mM) in PBS at r.t. for 1 h, followed by five washes with PBS. The cells were incubated with 5 μg mL^-1^ streptavidin-dye in PBS containing 1% (w/v) BSA for 1 h and washed three times with PBS.

For alkyne-Hoechst, U2OS cells were treated with 2 μM alkyne-Hoechst for 45 min and washed with warm DMEM for 15 min, and followed with the same procedures of DNA labeling experiments. Before gelation, alkyne-Hoechst-labeled cells were treated with 30 unit mL^-1^ DNase I for 30 min at 37°C to fragment the genomic DNA.

### Gelation, digestion, and expansion

Expansion microscopy was performed as described before with some modifications^3,4^. For anchoring with AcX, cells are incubated in AcX diluted in PBS at a concentration of 0.1 mg mL^-1^ overnight at r.t. followed by washing three times with PBS. For anchoring with GA, cells are incubated with 0.25% (v/v) GA in PBS for 10 min followed by washing three times with PBS. Monomer solution (1×PBS, 2 M NaCl, 2.5% (w/w) acrylamide, 0.15% (w/w) N,N’-methylenebisacrylamide, 8.625% (w/w) sodium acrylate) was mixed, frozen in aliquots, and thawed before use. Fresh prepared 10% (w/w) tetramethylethylenediamine (TEMED) and 10% (w/w) ammonium persulfate (APS) were diluted in monomer solution to final concentration of 0.2% (w/w) in gelation solution. Cells were incubated with the gelation solution at 4°C for 5 min, and then transferred to a humidified 37°C incubator for 1 h for gelation. The hydrogel was then digested in digestion buffer (50 mM Tris pH 8.0, 1 mM EDTA, 0.1% (v/v) Triton X-100, 0.8 M guanidine HCl) containing 8 unit mL^-1^ proteinase K at 37°C for different time, 4 h for AcX-treated samples and 2 h for GA-treated samples. The hydrogel was removed from digestion buffer and placed into deionized water to expand. Water was exchanged every 20 min until expansion was complete. If necessary, nucleus were stained with Hoechst 33342 to help locate cell position in the hydrogel.

### isHCR amplification

isHCR amplification was performed as described before with some modifications^7^. All reagents were dissolved in HCR amplification buffer (5× sodium chloride citrate (SSC) buffer, 0.1% (v/v) Tween-20, and 10% (w/v) dextran sulfate in ddH_2_O). Snap-cooled a pair of DNA-fluorophore HCR amplifiers separately in 5× SSC buffer by heating them at 95 °C for 90 s and then cooling them to r.t. over 30 min. Cells labeled with streptavidin were incubated with 0.5 µM DNA-biotin HCR initiators at r.t. for 1 h. Cells were then washed five times with PBS, and incubated with amplification buffer contained with 150 nM a pair of DNA-fluorophore HCR amplifiers overnight at r.t.. Cells were then washed five times with PBS. The following DNA sequences were used for isHCR amplification:

B1 Amplifier H1: 5’-AF546-CGTAAAGGAAGACTCTTCCCGTTTGCTGCCCTCCTCGCATTCTTTCTTGAGGAGGGCAGCAAACGGGAAGAG-3’

B1 Amplifier H2: 5’-GAGGAGGGCAGCAAACGGGAAGAGTCTTCCTTTACGCTCTTCCCGTTTGCTGCCCTCCTCAAGAAAGAATGC-AF546-3’

B1I2: 5’-Acrydite-ATATAGCATTCTTTCTTGAGGAGGGCAGCAAACGGGAAGAG-Biotin-3’

### Biotin trimer amplification

Cells labeled with streptavidin-dye were incubated with 50 µM biotin trimer at r.t. for 30 min, and washed five times with PBS. Cells were then incubated with 5 μg mL^-1^ streptavidin-dye in PBS containing 1% (w/v) BSA for 30 min, and washed five times with PBS. Desired amplified signal can be obtained by repeating this cycle.

### Imaging and data processing

All the expanded samples were performed imaging acquisition within 24 h. Expansion factor was quantified accurately by the hydrogel size of pre- and post-expanded gel air-water boundary. To minimize expanded samples drift during data acquisition, 24×50 mm rectangular #1.5 coverglasses (high-precision, Fisher Scientific) were coated with 0.1 mg mL^-1^ poly-D-lysine for 10 min at r.t. and allowed to dry. Excess water was carefully removed using laboratory wipes, and expanded samples could be stably attached to the glass surface and maintain immobility for a long time data collection. Confocal microscopy imaging was mainly performed on a commercial Leica SP8X laser scanning confocal system with a 63 ×/ 1.40-NA (numerical aperture) oil-immersion objective. White light laser (470-670 nm) was set to 70% of maximum power, and a 405 nm diode laser was used for Hoechst 33342 excitation. Some samples were imaged on a Zeiss LSM 700 laser scanning confocal system with a 20×/ 0.8-NA air objective (**Fig. 3b**), a 40×/ 1.40-NA oil-immersion objective (**Supplementary Fig. 4a**) and 63×/ 1.40-NA oil-immersion objective (**Fig. 2b,e** and **Fig. 3d**). The pinhole during all the image acquisition was opened at 1 Airy unit. The detector gain were optimized for each sample individually to achieve a desired level of intensity in the images collected before and after expansion. SIM imaging (**Fig. 3c**) was performed on a Nikon N-SIM microscopy system with a CFI SR Apo TIRF 100×/ 1.49-NA oil-immersion objective. 488 nm solid-state laser was used for excitation, and EMCCD (Andor DU-897) was used for collecting fluorescence signal. FIJI/ImageJ (https://fiji.sc, NIH) and Imaris (Bitplane) were used for processing, measurement, rendering and visualization of all the images in this study. Some images (**Fig. 2b** and **Supplementary Fig. 7e**) were deconvolved by Huygens software (Scientific Volume Imaging) using theoretical point spread functions and water as ‘Mounting Medium’.

### Data availability

The data and probes in this study are available from the corresponding author upon reasonable request.

