## Supplemental Material for "Click-ExM enables expansion microscopy for all biomolecules"

### SUPPLEMENTARY INFORMATION FILE

#### Supplementary Figures 1-13

##### Click-ExM enables expansion microscopy for all biomolecules

De-en Sun[1,2], Xinqi Fan[1,2], Hao Zhang[1,2], Zhimin Huang[1,3], Qi Tang[1,2], Wei Li[1,2], Jinyi Bai[1,3], Xiaoguang Lei[1,2,3,4,5], Xing Chen[1,2,3,4,5]*

[1]College of Chemistry and Molecular Engineering, [2]Beijing National Laboratory for Molecular Sciences, [3]Peking-Tsinghua Center for Life Sciences, [4]Synthetic and Functional Biomolecules Center, [5]Key Laboratory of Bioorganic Chemistry and Molecular Engineering of Ministry of Education, Peking University, Beijing 100871, China.

*

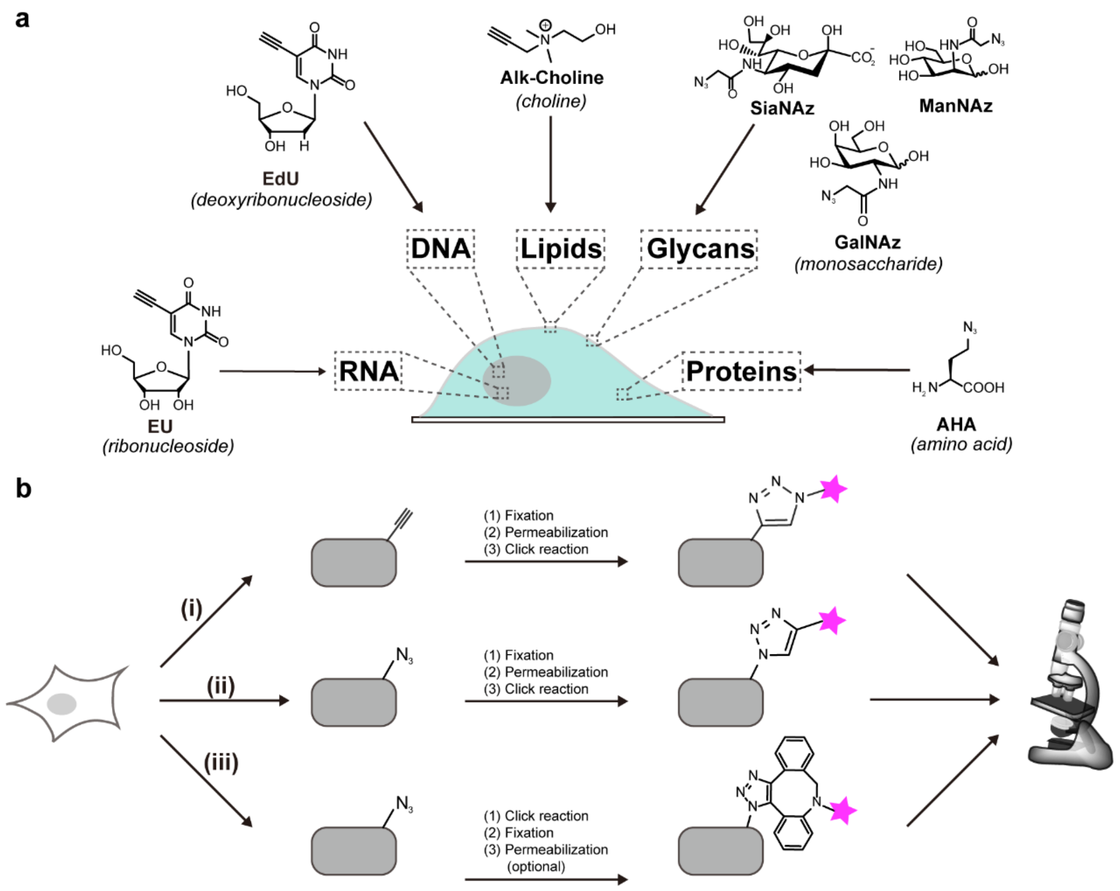


**Supplementary Fig. 1**

**Click labeling enables tagging of various biomolecules with fluorophores.**

(**a**) The building blocks of various biomolecules including RNA, DNA, lipids, glycans and proteins are derivatized with a biorthogonal or "clickable" functional group (e.g., the azide or alkyne). The clickable monomers are metabolically incorporated into the corresponding biomolecules. (**b**) The alkyne-incorporated biomolecules are reacted with azide-fluorophore via CuAAC (i); the azide-incorporated biomolecules are reacted with alkyne-fluorophore via CuAAC (ii) or with dibenzocyclooctyne-fluorophore (DBCO-fluorophore) via copper-free click chemistry (iii). The fluorescently labeled biomolecules are visualized by conventional fluorescence microscopy.

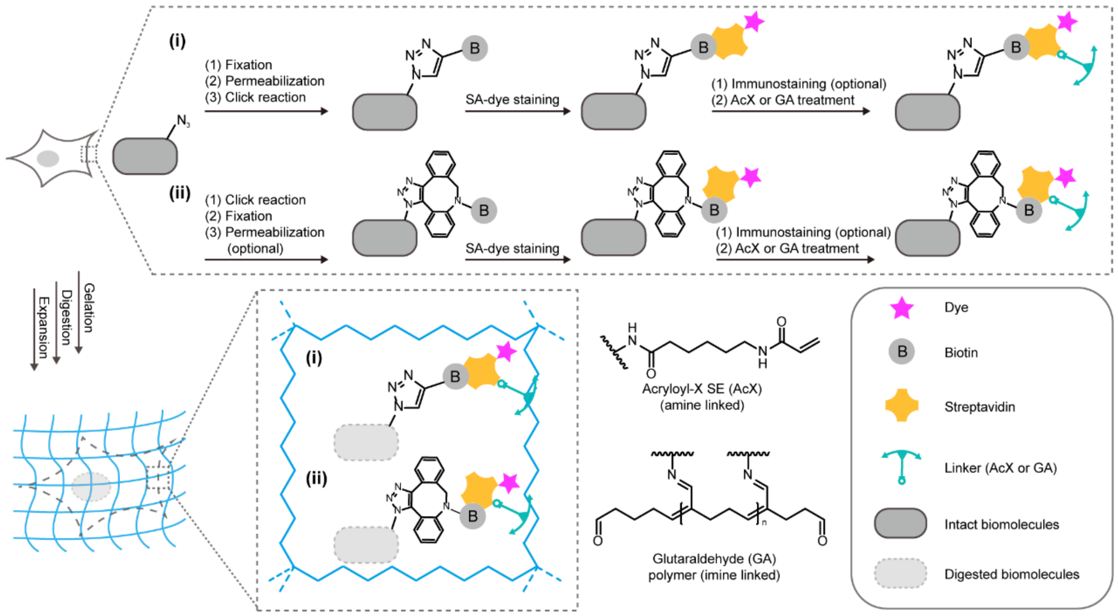


**Supplementary Figure 2 (Related to Fig. 1a)**

**Schematic of the click-ExM workflow for cells metabolically labeled with azides.**

After metabolic labeling of biomolecules with azides, the cells are reacted with alkyne-biotin via click chemistry (i) or DBCO-biotin via copper-free click chemistry (ii) and staining with streptavidin-dye (SA-dye), followed with the standard ExM procedure including steps of AcX or GA treatment, gelation, digestion and expansion.

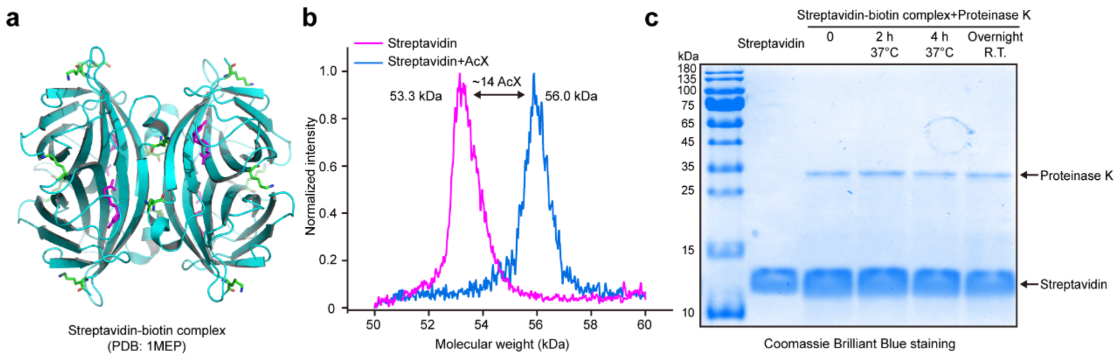


#### Supplementary Figure 3

**Streptavidin provides multiple anchoring sites and resists proteinase K digestion.**

(**a**) Crystal structure of the streptavidin-biotin complex (PDB code 1MEP). Streptavidin, biotin and lysine residues are shown in cyan, magenta, and green, respectively. (**b**) MALDI-TOF spectra of streptavidin (magenta) and AcX-conjugated streptavidin (blue). Corresponding mass shift indicates a total of ~14 AcX molecules conjugated with streptavidin. (**c**) SDS-PAGE gel showing streptavidin (100 μg) treated with proteinase K (8 unit $mL^{-1}$) in ExM digestion buffer containing 600 μM biotin for 0 h, 2 h and 4 h at 37 °C, or overnight at r.t. Untreated streptavidin was included as a control. All experiments were independently performed ≥3 times except the MALDI-TOF MS experiment in **b** (one experiment); representative data are shown.

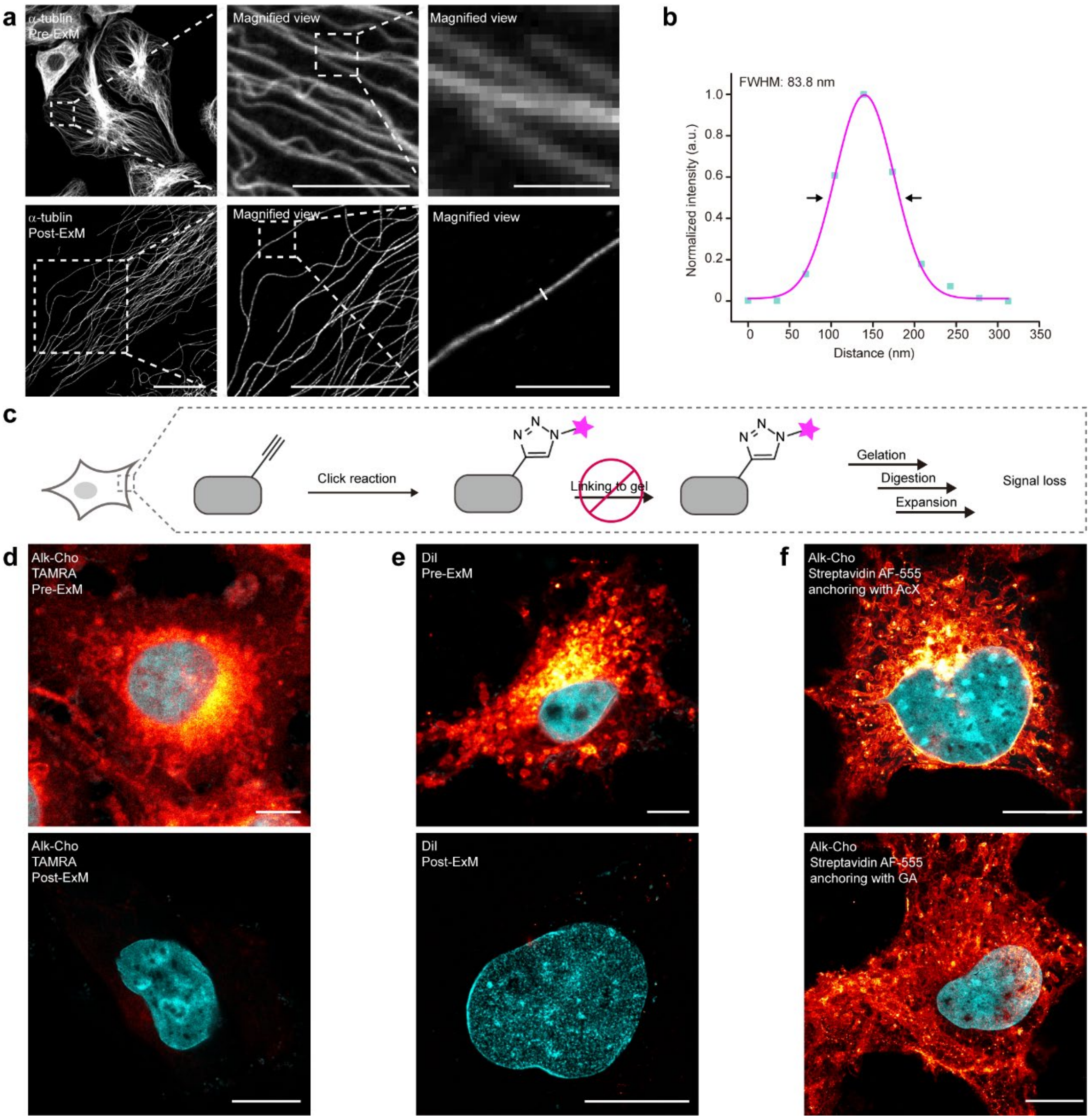


**Supplementary Figure 4**

**Click-ExM preserves lipid labeling signal by streptavidin-fluorophore conjugates.**

(**a**) Protein-retention ExM protocol adapted for click-ExM. Pre- and post-ExM images of immunostained α-tubulin (AF555, gray). (**b**) Gaussian fitting plot of the intensity profile along the line in **a**, in which full-width-half-maximum (FWHM) was calculated as 83.8 nm after rescaling by the expansion ratio. (**c**) The cells were treated with alk-Cho to metabolically label cho-containing phospholipids, followed by reaction with azide-dye and the standard ExM procedure. (**d**) Pre- and post-ExM images of COS-7 cells treated with alk-Cho and reacted with alkyne-TAMRA (red hot). (**e**) Pre- and post-ExM images of COS-7 cells stained with the lipophilic dye, 1,1′-dioctadecyl-3,3,3′,3′-tetramethylindocarbocyanine perchlorate (DiI, red hot). (**f**) Click-ExM imaging of COS-7 cells treated alk-Cho, reacted azide-biotin, and stained with streptavidin-AF555 (red hot) was compatible with anchoring with AcX (top) and GA (bottom). Nucleus were stained by Hoechst 33342 (cyan). AcX was used for anchoring unless otherwise specified. Scale bars: 10 μm (**a**, left and middle, **d**-**f**) and 2 μm (**a**, right). All distances and scale bars are corresponding to the pre-expansion dimension. All images were taken. All experiments were independently performed with a confocal microscope ≥3 times except the DiI staining experiment in **e** (two independent experiments); representative data are shown.

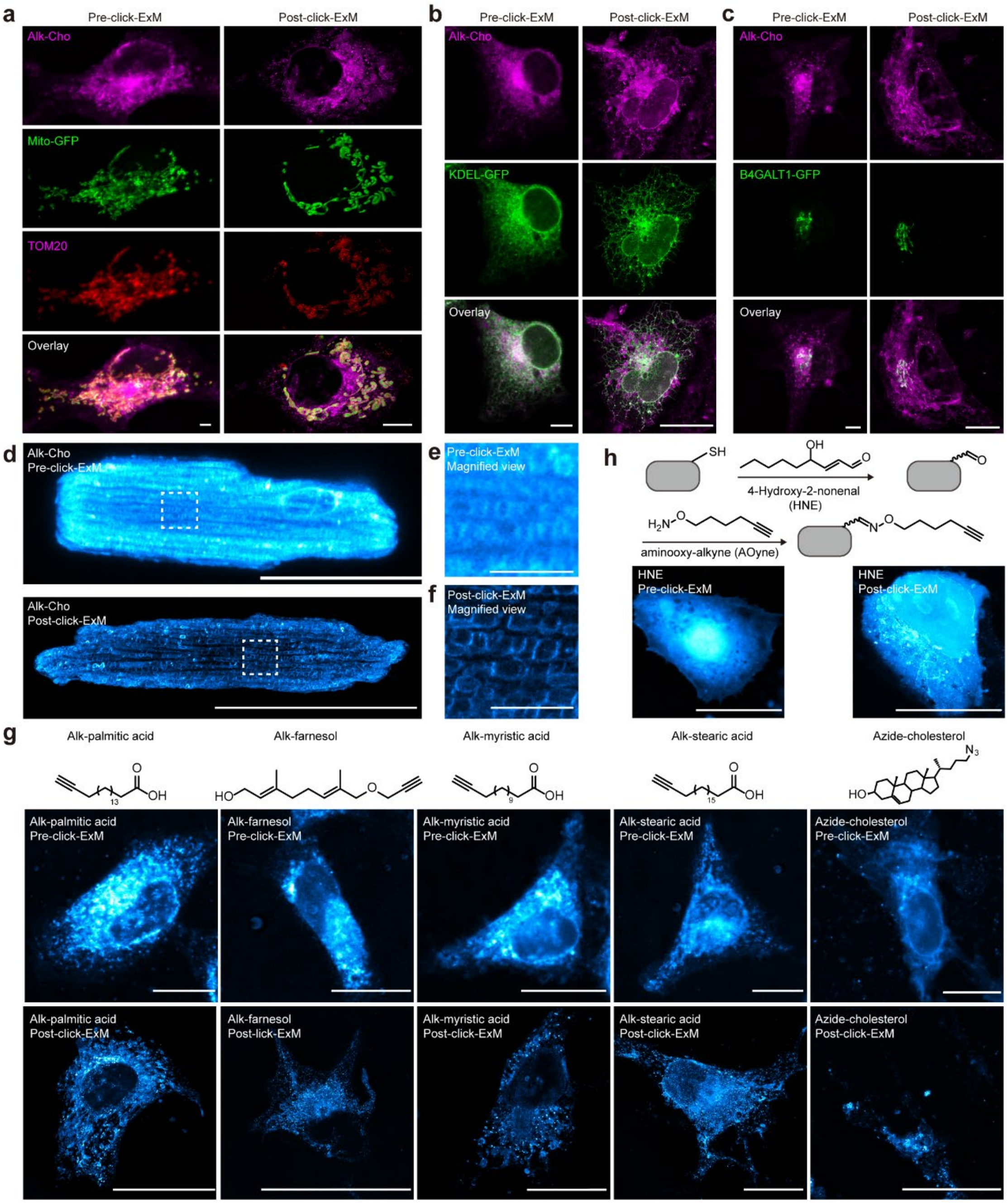


**Supplementary Figure 5 (Related to Fig. 1b-h)**

**Pre- and post-click-ExM images of lipids.**

(**a**) Pre- and post-click-ExM images of alk-Cho-labeled phospholipids (AF555, magenta) in COS-7 cells. Mito-GFP (green) was expressed in the mitochondrial matrix and TOM20 located in the mitochondrial outer membrane was immunostained with AF647 (red). (**b**) Pre- and post-click-ExM images of alk-Cho-labeled phospholipids (AF555, magenta) and ER expressing KDEL-GFP (green) in COS-7 cells. (**c**) Pre- and post-click-ExM images of alk-Cho-labeled phospholipids (AF555, magenta) in COS-7 cells with the Golgi expressing B4GALT1-GFP (green). The same post-click-ExM images were shown in **fig. 1**. (**d**) Pre- and post-click-ExM images of alk-Cho-labeled phospholipids (AF555, cyan hot) in rat cardiomyocytes. (**e**) Magnified views of the boxed region from the pre-click-ExM image in **d**. (**f**) Magnified views of the boxed region from the post-click-ExM image in **d**. (**g**) Pre- and post-click-ExM images of lipids in COS-7 cells metabolically labeled with various bioorthgonal reporters including alkynyl palmitic acid, alkynyl-farnesol, alkynyl myristic acid, alkynyl stearic acid and azide-cholesterol (AF555, cyan hot). (**h**) Pre- and post-click-ExM images of 4-hydroxy-2-nonenal (HNE)-labeled protein carbonylation (AF555, cyan hot) in COS-7 cells. Cysteine residues were reacted with HNE through Michael addition, reacted with aminooxy-alkyne (AOyne), and followed with the click-ExM workflow. Scale bars: 50 μm (**d**, **g**, **h**), 10 μm (**a-c**) and 5 μm (**e**, **f**). Scale bars are corresponding to the pre-expansion dimension. GA (**a-d**) and AcX (**g**) were used for anchoring. All experiments were independently performed ≥3 times with a confocal microscope.

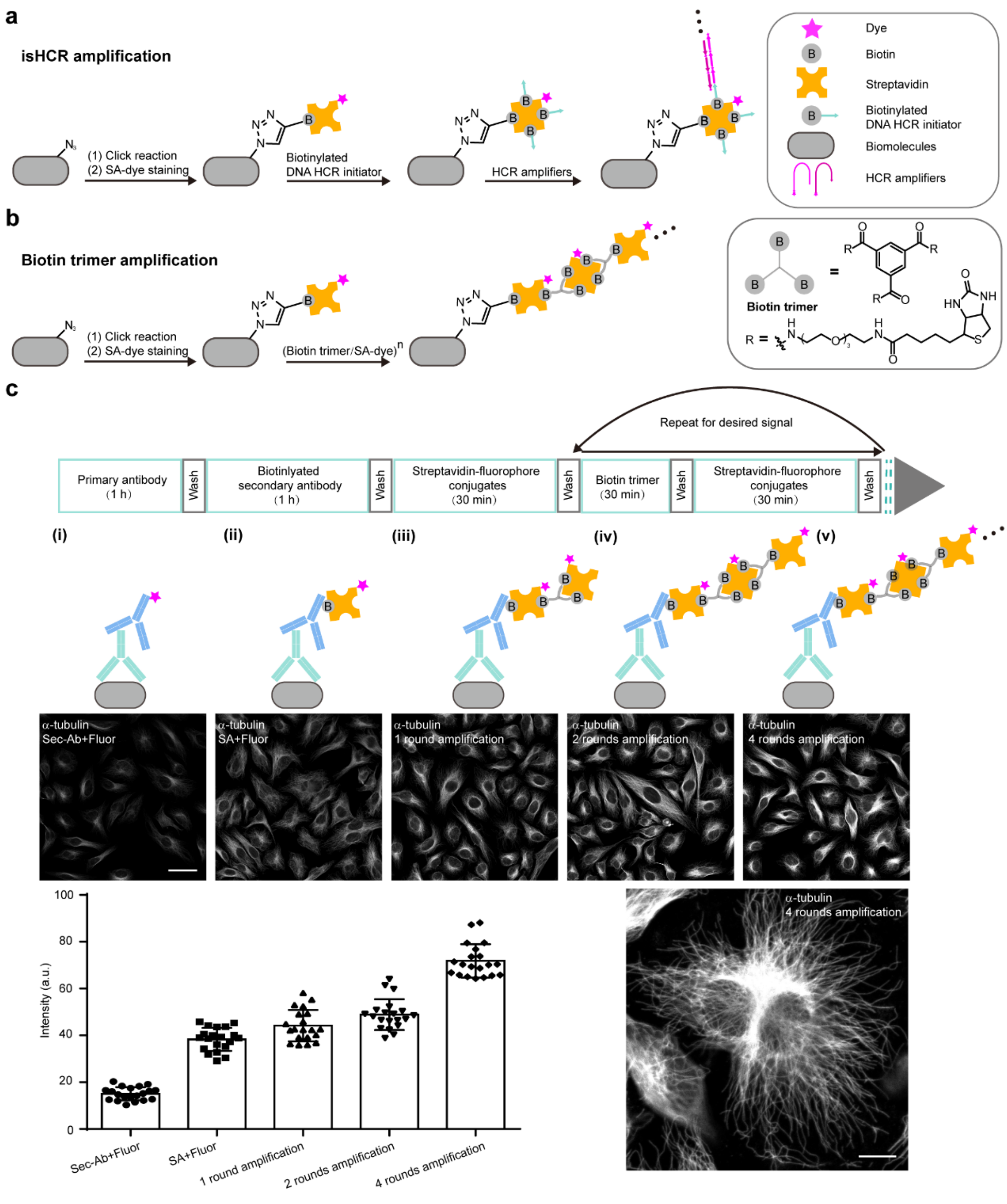


#### Supplementary Figure 6

**Signal amplification by immunosignal hybridization chain reaction (isHCR) or by a biotin trimer.**

(**a**) Workflow of isHCR for signal amplification. Cells click-labeled with streptavidin-fluorophore conjugates (SA-dye) were incubated with DNA–biotin HCR initiators, and then incubated with a pair of DNA–fluorophore HCR amplifiers for signal amplification. (**b**) Workflow of iterative signal amplification by using the biotin trimer. (**c**) Fluorescence imaging of HeLa cells with α-tubulin immunostained with AF555, which was amplified by using the biotin trimer. Procedures of iterative signal amplification (top). HeLa cells were immunostained for α-tubulin, and five labeling conditions were used for comparison from left to right: (i) conventional antibody staining with AF555-conjugated secondary antibody, (ii) biotin-conjugated secondary antibody and streptavidin AF555, (iii)-(v) one, two and four rounds of iterative staining by biotin trimer and streptavidin-AF555. Representative images of each conditions were shown (middle). Histogram shows fluorescence intensity quantification of different cells in five labeling conditions (bottom left). Bars represent the mean value and error bars represent the standard deviation. Magnified image for four rounds signal amplification of immunostained α-tubulin in HeLa cells was shown (bottom right). Scale bars: 50 μm (**c**, middle) and 10 μm (**c**, bottom right). All experiments were independently performed ≥3 times with a confocal microscope.

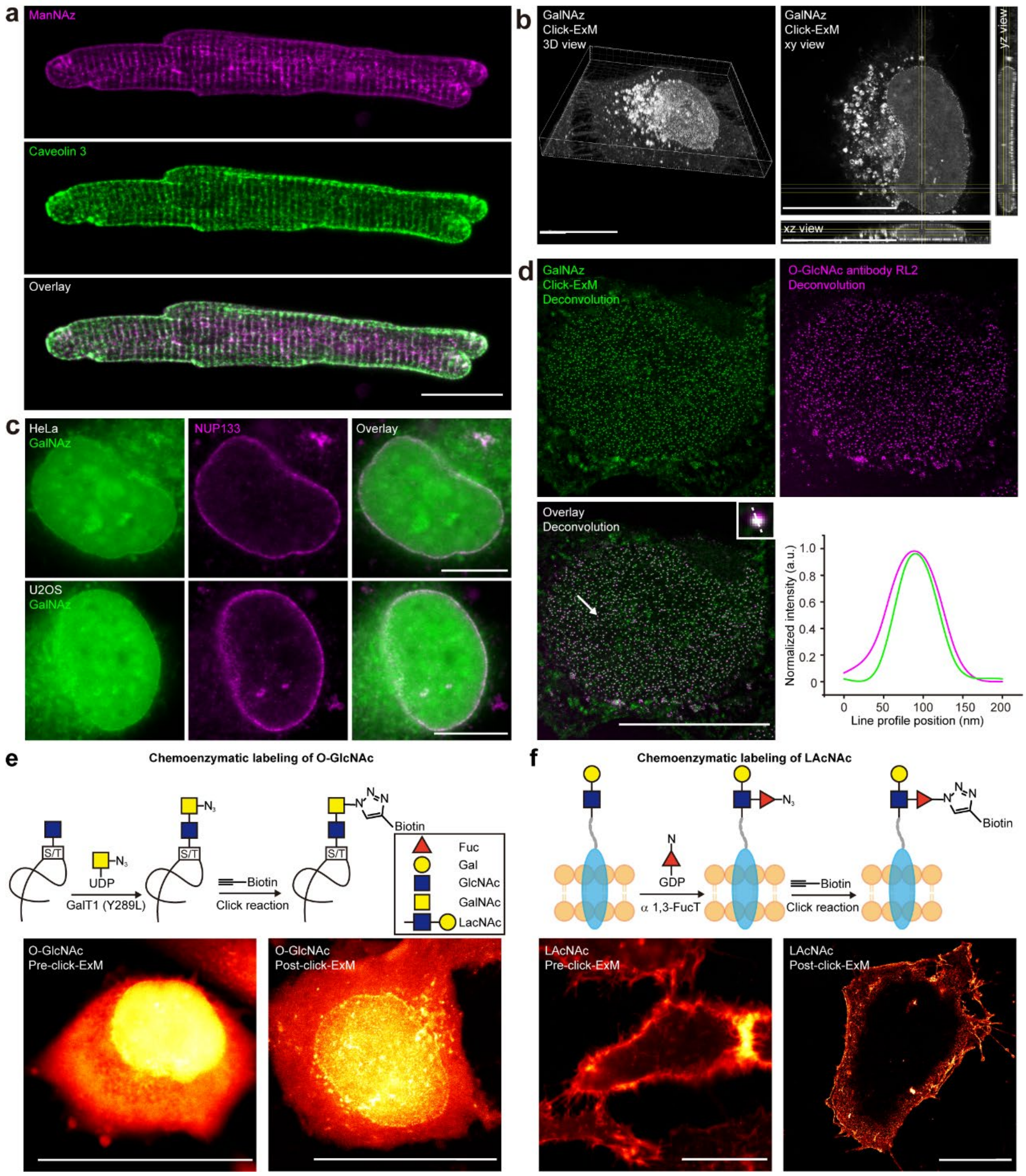


**Supplementary Figure 7 (Related to Fig. 2)**

**Pre- and post-click-ExM imaging of glycans.**

(**a**) Confocal fluorescence images of cardiomyocyte, in which sialoglycans were metabolically labeled with ManNAz (AF555, magenta) and the T-tubule network was immunostained by using the T-tubule marker caveolin 3 (AF488, green). (**b**) Click-ExM images of GalNAz-labeled HeLa cells in **Fig.2e** shown in 3D (left) and xy, xz, yx views (right). 3D images were rendered in Imaris using the MIP 3D mode. (**c**) Colocalization between GalNAz-labeled O-GlcNAc (AF488, green) and immunostained NUP133 (AF555, magenta) in HeLa cells and U2OS cells. (**d**) Click-ExM images of O-GlcNAc in HeLa cells after deconvolution. O-GlcNAc was metabolically labeled by GalNAz (AF488, green), and immunostained with RL2 (AF546, magenta). Fluorescence intensity profiles of GalNAz (green) and RL2 (magenta) along the dotted lines in the boxed region were shown. (**e**) Pre- and post-click-ExM images of O-GlcNAc distribution chemoenzymatically labeled by Y289L GalT in CHO cells (AF555, red hot). (**f**) Pre- and post-click-ExM images of LacNAc-containing glycans (AF555, red hot), which were chemoenzymatically labeled by α1,3-FucT in HeLa cells. Scale bars: 20 μm (**a**, **c**, **e**, **f**) and 10 μm (**b**, **d**). All distances and scale bars are corresponding to the pre-expansion dimension. AcX was used for anchoring. All experiments were independently performed ≥3 times with a confocal microscope.

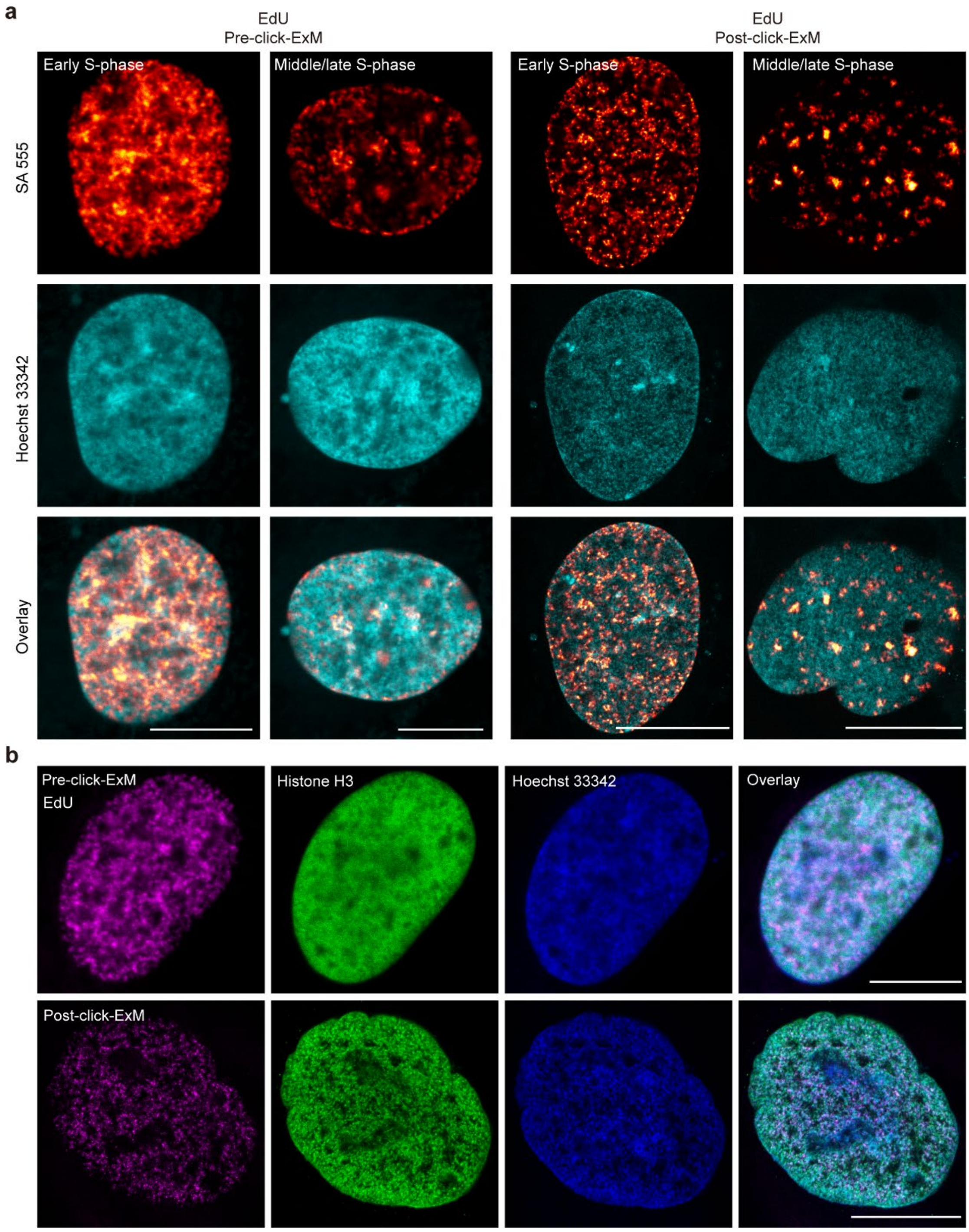


**Supplementary Figure 8 (Related to Fig. 3c)**

**Pre- and post-click-ExM imaging of EdU-labeled nascent DNA in U2OS cells.**

(**a**) Pre- and post-click-ExM images of EdU-labeled nascent DNA (AF555, red hot) in early and middle/late S-phase, from which chromatins with different sizes and shapes were observed. Nucleus were stained with Hoechst 33342 (cyan). (**b**) Click-ExM images showing co-localization of nascent DNA (AF555, magenta) with histone H3

(AF488, green). Nucleus were stained with Hoechst 33342 (blue). Scale bars: 10 μm (**a**, **b**). Scale bars are corresponding to the pre-expansion dimension. AcX was used for anchoring. All experiments were independently performed with a confocal microscope ≥3 times except for histone H3 immunostaining in **b** (two independent experiments).

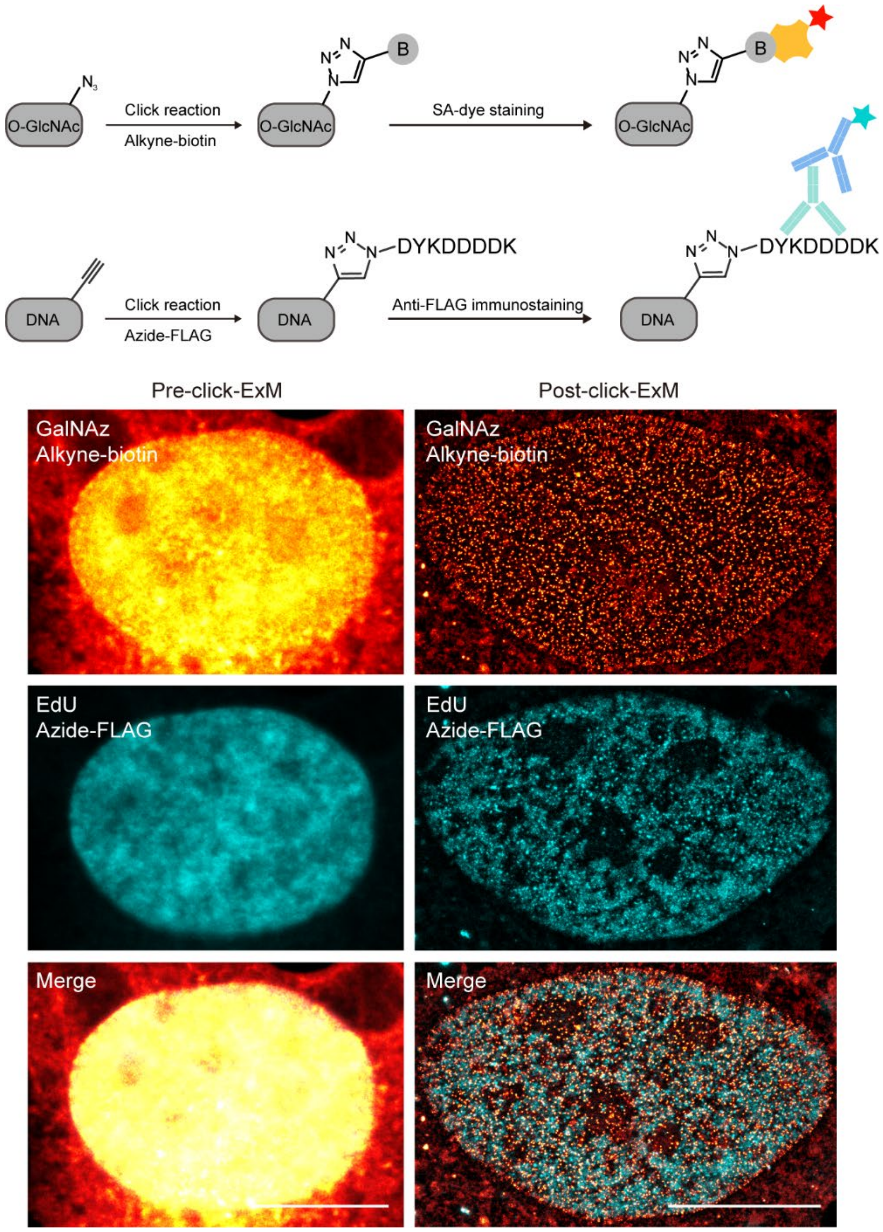


**Supplementary Figure 9**

**Two-color click-ExM imaging of O-GlcNAc and DNA.**

HeLa cells were simultaneously treated with GalNAz and EdU. The azide-incorporated O-GlcNAc was reacted with alkyne-biotin and stained with streptavidin-AF488 (red hot). The alkyne-incorporated nascent DNA was reacted with azide-FLAG and immunostained with FLAG antibody (AF555, cyan). Pre- and post-click-ExM images are shown. AcX was used for anchoring. Scale bars: 10 μm. All scale bars are corresponding to the pre-expansion dimension. The experiment was independently performed 3 times with a confocal microscope.

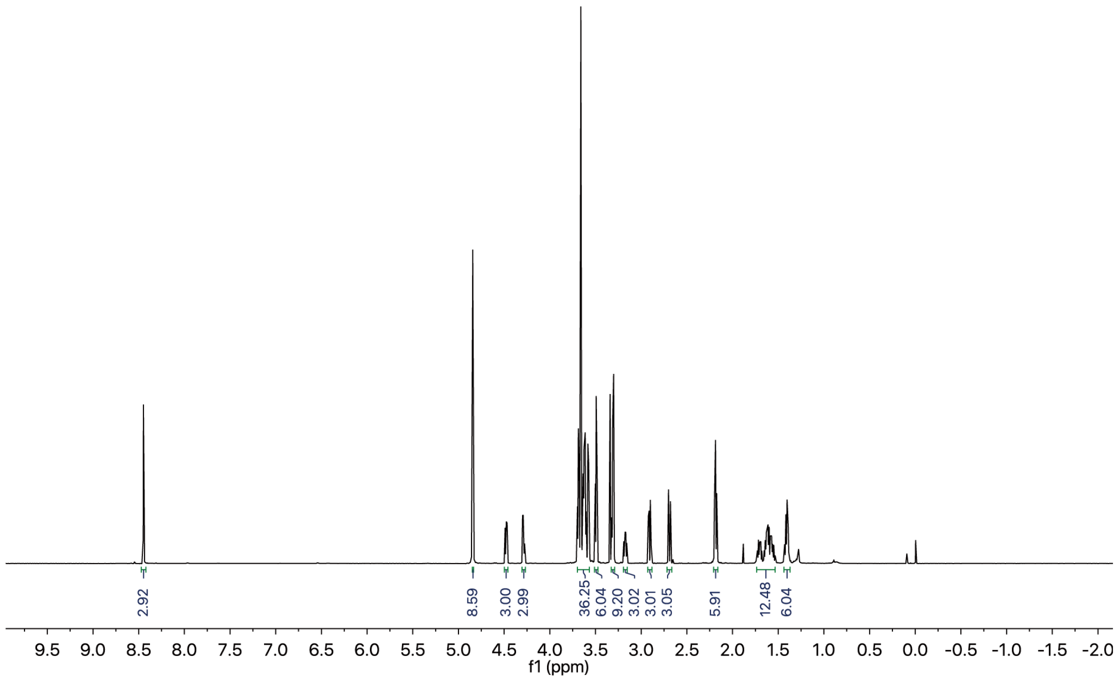


**Supplementary Figure 10**

**$^{1}$H-NMR spectrum of biotin trimer.**

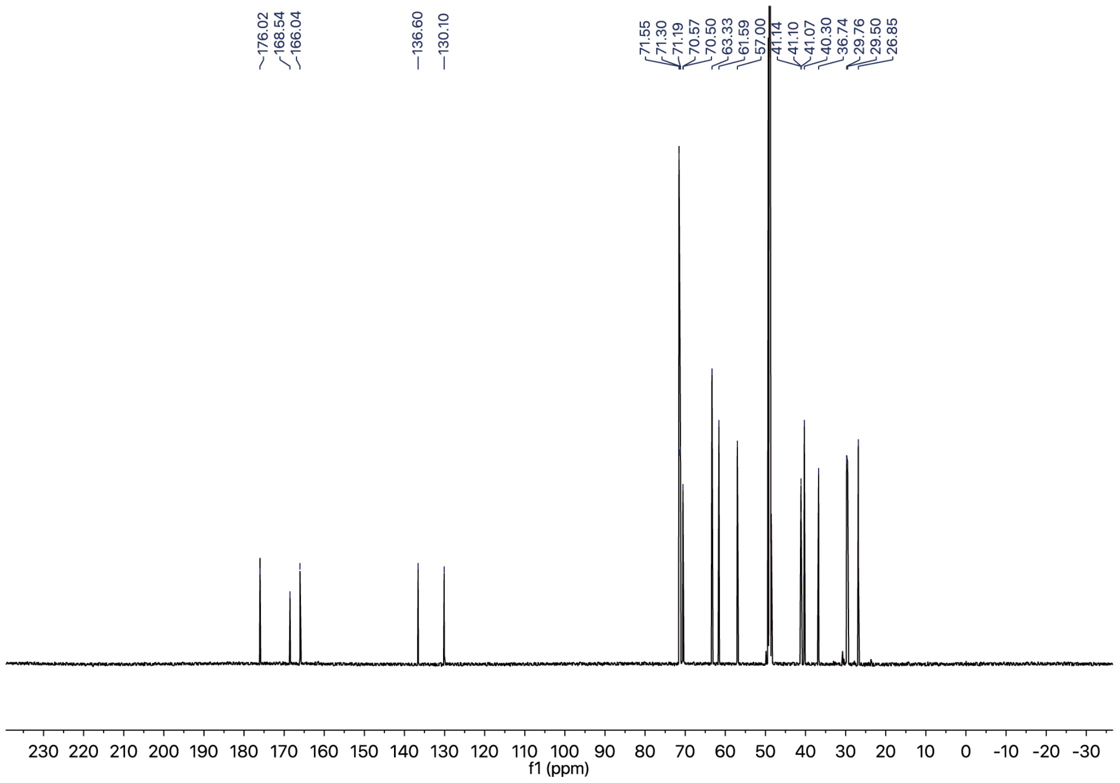


**Supplementary Figure 11**

**$^{13}$C-NMR spectrum of biotin trimer.**

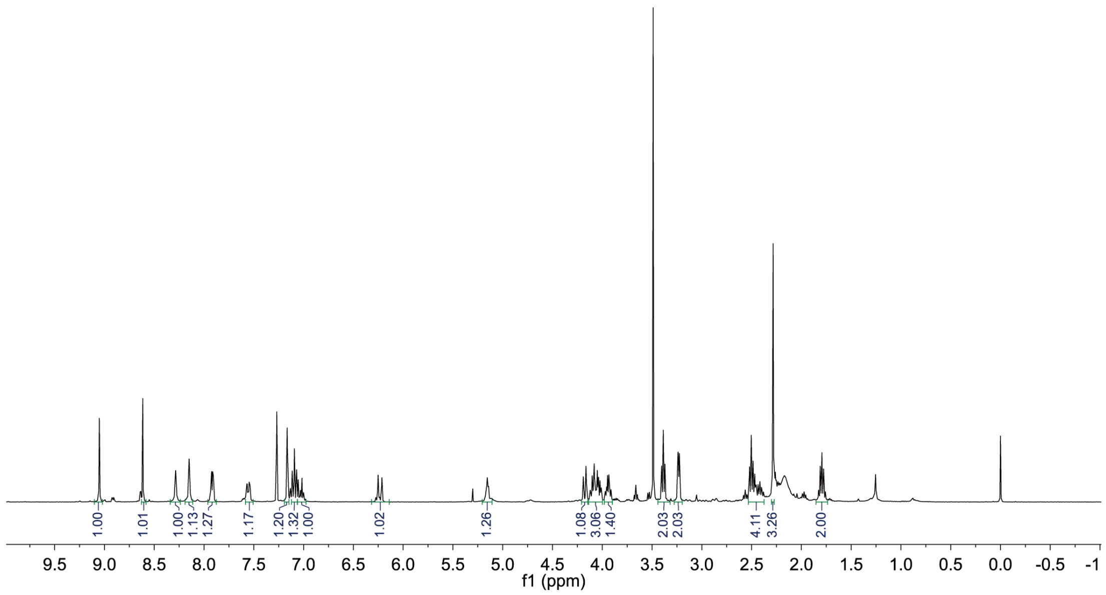


**Supplementary Figure 12**

**$^1$H-NMR spectrum of azide-afatinib.**

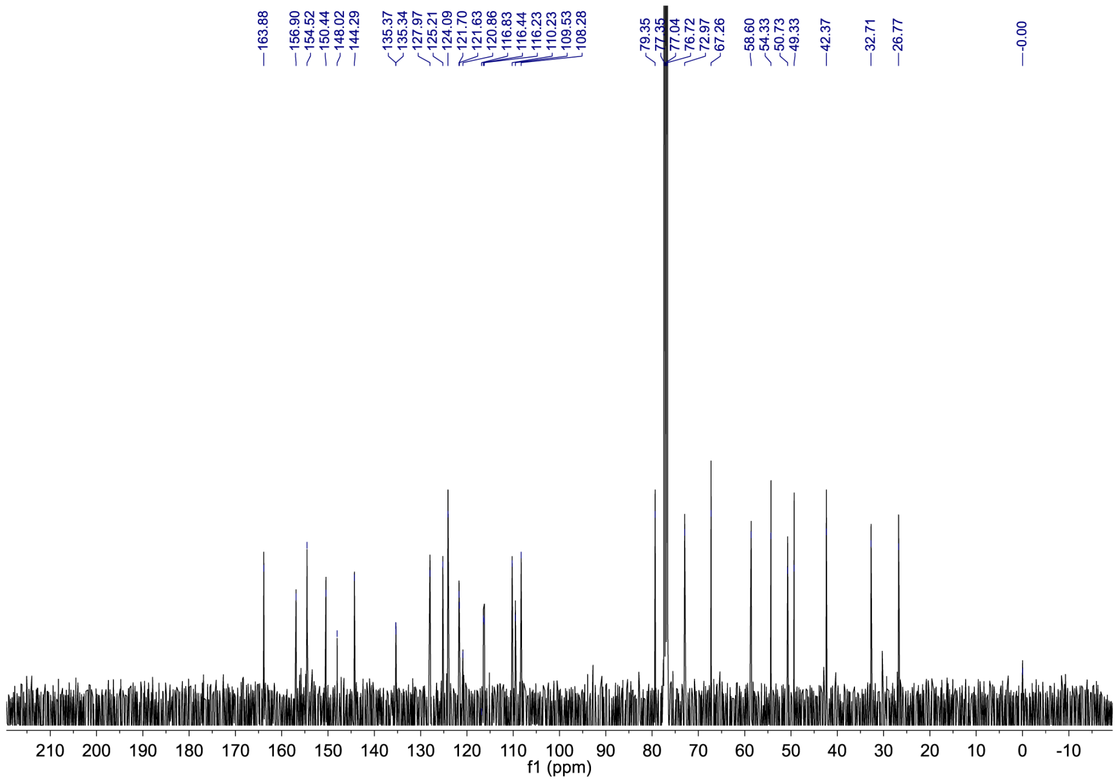


**Supplementary Figure 13**

**$^{13}$C-NMR spectrum of azide-afatinib.**

### Supplementary Note 1

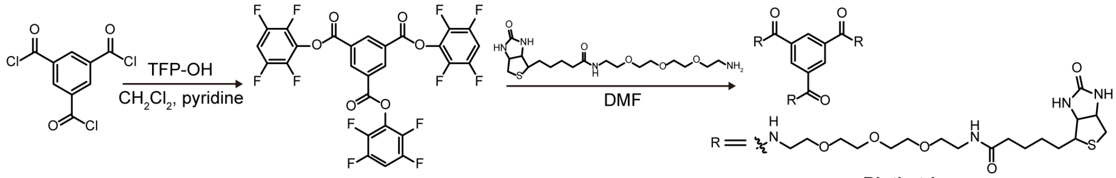


**Biotin trimer**

**Synthesis of biotin trimer.** To a solution of biotin-PEG$_3$-amine (10 mg) in 100 μL dimethyl formamide (DMF), 100 μL 1,3,5-benzene tricarboxylic acid tris-(2,3,5,6-tetrafluorophenyl) ester (0.474 mg mL$^{-1}$ in DMF) was added at r.t. The reaction mixture was stirred at r.t. overnight followed by removal of solvent under vacuum. The crude product was further purified by flash chromatography with methanol (MeOH): ethyl acetate (EtOAc) gradually from the volume ratio at 1 : 100 to 3 : 7 to give the final compound as white solid (7 mg, 68%).

$^1$H NMR (500 MHz, $CD_3OD$) 8.45 (s, 3H), 4.84 (s, 9H), 4.50-4.46 (m, 3H), 4.30-4.27 (m, 3H), 3.70-3.56 (m, 36H), 3.51-3.48 (m, 6H), 3.33-3.29 (m, 9H), 3.19-3.15 (m, 3H), 2.93-2.88 (m, 3H), 2.69 (d, *J*=10.5 Hz, 3H), 2.21-2.16 (m, 6H), 1.74-1.53 (m, 12H), 1.44-1.37 (m, 6H).

$^{13}$C NMR (125 MHz, $CD_3OD$) δ 176.02, 168.54, 166.04, 136.60, 130.10, 71.55, 71.30, 71.19, 70.57, 70.50, 63.33, 61.59, 57.00, 41.14, 41.10, 41.07, 40.30, 36.74, 29.76, 29.50, 26.85.

HRMS (ESI): m/z calculated for $C_{63}H_{103}N_{12}O_{18}S_3$ $[M+H]^+$ 1411.6670, found 1411.6634.

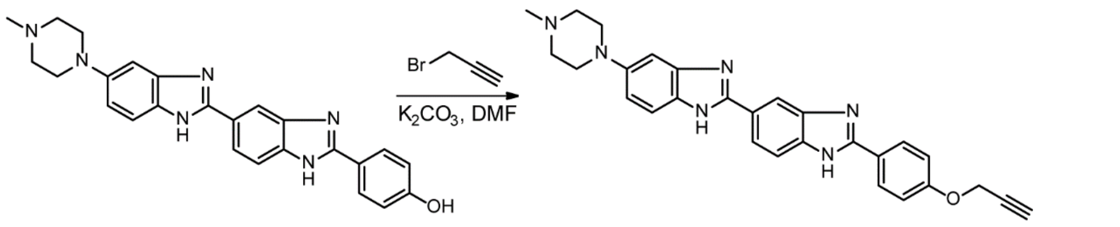


**Alkyne-Hoechst**

**Synthesis of alkyne-Hoechst.** To a stirred solution containing DMF (5 mL), Hoechst 33258·3HCl (50 mg, 0.094 mmol) and potassium carbonate (80 mg, 0.58 mmol), propargyl bromide (80% (v/v) in toluene, 16 μL, 0.14 mmol) was added under nitrogen atmosphere. The reaction mixture was stirring for 12 h in darkness at r.t., followed by removal of solvent under vacuum. The crude product was further purified by preparative high performance liquid chromatography (HPLC), acidified by hydrochloric acid and lyophilized to give product as a yellow powder (41 mg, 76% yield).

HRMS (ESI): m/z calculated for $C_{28}H_{27}N_6O$ $[M+H]^+$ 463.2241, found 463.2241.

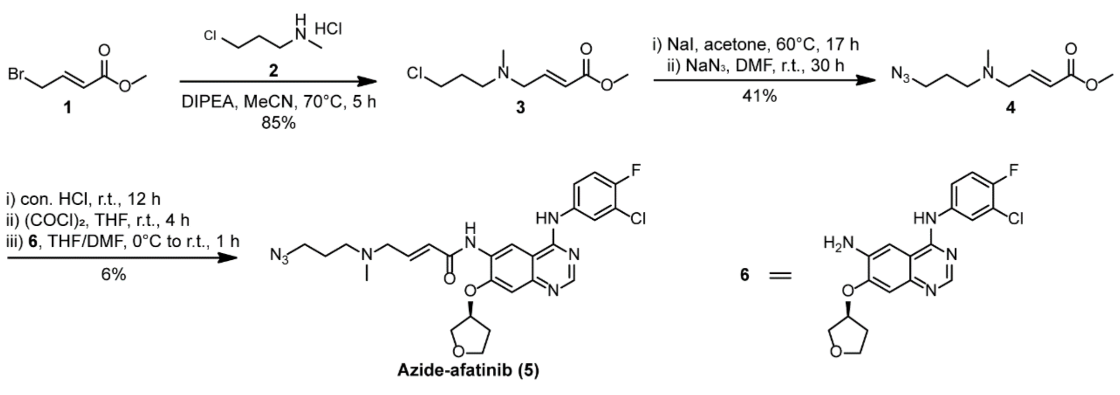


**Azide-afatinib (5)**

**Synthesis of azide-afatinib.**

Methyl (*E*)-4-((3-chloropropyl)(methyl)amino)but-2-enoate (**3**)

To a solution of **2** (0.16 g, 1.1 mmol) in acetonitrile (MeCN, 8 mL), **1** (0.18 g, 1.0 mmol) was added, after which N,N-diisopropylethylamine (DIPEA, 0.54 mL, 3.1 mmol) was added dropwise. The reaction mixture was heated at 70°C for 5 h. After cooling to r.t., the resulting mixture was concentrated *in vacuo* to remove the solvent, redissolved with EtOAc (100 mL), and then washed with sat. $NaHCO_3$ and brine. The EtOAc phase was dried over anhydrous $Na_2SO_4$, and the filtrate was concentrated *in vacuo*. The residue was subjected to a column of silica gel (1%~2% MeOH in Dichloromethane (DCM)) to afford compound **3** (0.18 g, 85%) as a light brown oil.

$^1$H NMR (400 MHz, $CDCl_3$) δ 6.95 (dt, $J_1$ = 15.7 Hz, $J_2$ = 6.1 Hz, 1H), 5.99 (dt, $J_1$ = 15.7 Hz, $J_2$ = 1.5 Hz, 1H), 3.74 (s, 3H), 3.60 (t, $J$ = 6.5 Hz, 2H), 3.15 (dd, $J_1$ = 6.0 Hz, $J_2$ = 1.5 Hz, 2H), 2.51 (t, $J$ = 6.8 Hz, 2H), 2.23 (s, 3H), 1.92 (m, 2H); ESI-MS: $[M+H]^+$ 206.2.

Methyl (*E*)-4-((3-azidopropyl)(methyl)amino)but-2-enoate (**4**)

To a solution of **3** (0.13 g, 0.63 mmol) in acetone (2 mL), NaI (0.29 g, 1.9 mmol) was added. The reaction mixture was heated at 60°C for 17 h. Crude NMR showed the complete conversion of **3** to iodo-intermediate. After cooling to r.t., the solvent was removed *in vacuo*. The residue was redissolved with dry DMF (2 mL), to which $NaN_3$ (0.21 g, 3.2 mmol) was added in one portion. The reaction mixture was stirred at r.t. for 30 h. Crude NMR showed the complete conversion of iodo-intermediate to the product **4**. EtOAc (60 mL) was added to the mixture, and the resulting organic phase was washed with sat. $NaHCO_3$ and brine. The organic phase was dried over anhydrous $Na_2SO_4$ and concentrated *in vacuo*. The residue was subjected to a column of silica gel (Petroleum ether (PE)/EtOAc = 2:1 to 1:1) to afford **4** (55 mg, 0.26 mmol, 41%) as a light yellow oil.

$^1$H NMR (400 MHz, $CDCl_3$) δ 6.94 (dt, $J_1$ = 15.7 Hz, $J_2$ = 6.1 Hz, 1H), 5.99 (d, $J$ = 15.7 Hz, 1H), 3.75 (s, 3H), 3.35 (t, $J$ = 6.7 Hz, 2H), 3.14 (d, $J$ = 6.1 Hz, 2H), 2.45 (t, $J$ = 6.9 Hz, 2H), 2.22 (s, 3H), 1.74 (m, 2H); ESI-MS: $[M+H]^+$ 213.2.

Azide-afatinib (**5**)

Concentrated HCl (1.3 mL) was added to **4** (55 mg, 0.26 mmol). The mixture was stirred at r.t. for 12 h. The resulting mixture was concentrated *in vacuo* to remove most water and HCl. The residue was redissolved with ethanol (EtOH) and concentrated *in vacuo* to remove residual water. The crude intermediate was dried in high vacuo and redissolved with dry tetrahydrofuran (THF), to which oxalyl chloride (0.75 mL, 7.9 mmol) was added dropwise. The mixture was stirred at r.t. for 4 h. The excessive oxalyl chloride was removed *in vacuo* and the resulting residue was dried in high vacuo for 20 min. The residue was redissolved with dry THF (0.5 mL) and cooled to 0°C, to which the solution of **6** (94 mg, 0.25 mmol) in dry DMF (0.5 mL) was added dropwise. The reaction mixture was slowly warmed to r.t. in 1 h and quenched with one drop ammonia water. EtOAc (50 mL) was added to the mixture, and the resulting organic phase was washed with sat. $NaHCO_3$ and brine. The organic phase was dried over anhydrous $Na_2SO_4$ and concentrated *in vacuo*. The residue was subjected to a column of silica gel (MeOH in DCM = 1% ~ 3%) to afford **5** (9 mg, 0.06 mmol, 6%).

$^1$H NMR (400 MHz, $CDCl_3$) δ 9.04 (s, 1H), 8.61 (s, 1H), 8.27 (s, 1H), 8.14 (s, 1H), 7.91 (dd, $J_1$ = 6.6 Hz, $J_2$ = 2.6 Hz, 1H), 7.55 (m, 1H), 7.15 (s, 1H), 7.13-6.97 (m, 2H), 6.22 (d, $J$ = 15.2

Hz, 1H), 5.14 (t, $J = 5.0$ Hz, 1H), 4.17 (d, $J = 10.7$ Hz, 1H), 4.12-3.99 (m, 3H), 3.93 (m, 1H), 3.38 (t, $J = 6.6$ Hz, 2H), 3.22 (d, $J = 5.7$ Hz, 2H), 2.52-2.37 (m, 4H), 2.27 (s, 3H), 1.78 (m, 2H); $^{13}$C NMR (101 MHz, $CDCl_3$) δ 163.9, 156.9, 154.5, 150.4, 148.0, 144.3, 135.3, 128.0, 125.2, 124.1, 121.6, 120.9, 116.4, 116.2, 110.2, 109.5, 108.3, 79.3, 73.0, 67.3, 58.6, 54.3, 49.3, 42.4, 32.7, 26.8.

HRMS (ESI): m/z calculated for $C_{26}H_{29}ClFN_8O_3$, $[M+H]^+$ 555.2030, found 555.2018.
